# Over-expression of wild-type *ACVR1* in fibrodysplasia ossificans progressiva mice rescues perinatal lethality and inhibits heterotopic ossification

**DOI:** 10.1101/2021.12.08.471385

**Authors:** Masakazu Yamamoto, Sean J. Stoessel, Shoko Yamamoto, David J. Goldhamer

**Author notes:** **One Sentence Summary:** Over-expression of wild-type *ACVR1* inhibits heterotopic ossification in FOP mice.

## Abstract

Fibrodysplasia ossificans progressiva (FOP) is a devastating disease of progressive heterotopic bone formation for which effective treatments are currently unavailable. FOP is caused by dominant gain-of-function mutations in the receptor ACVR1 (also known as ALK2), which render the receptor inappropriately responsive to activin ligands. In previous studies, we developed a genetic mouse model of FOP that recapitulates most clinical aspects of the disease. In this model, genetic loss of the wild-type *Acvr1* allele profoundly exacerbated heterotopic ossification, suggesting the hypothesis that the stoichiometry of wild-type and mutant receptors dictates disease severity. Here, we tested this model by producing FOP mice that conditionally over-express human wild-type ACVR1. Injury-induced heterotopic ossification (HO) was completely blocked in FOP mice when expression of both the mutant and wild-type receptor were targeted to Tie2-positive cells, which includes fibro/adipogenic progenitors (FAPs). Perinatal lethality of *Acvr1^R206H/+^* mice was rescued by constitutive *ACVR1* over-expression and these mice survived to adulthood at predicted Mendelian frequencies. Constitutive over-expression of *ACVR1* also provided protection from spontaneous HO, and the incidence and severity of injury-induced HO in these mice was dramatically reduced. Analysis of pSMAD1/5/8 signaling both in cultured cells and *in vivo* indicates that *ACVR1* over-expression functions cell-autonomously by reducing osteogenic signaling in response to activin A. Manipulating the stoichiometry of FOP-causing and wild-type ACVR1 receptors may provide the foundation for novel therapeutic strategies to treat this devastating disease.

## Introduction

Heterotopic ossification (HO), the formation of bone at extraskeletal anatomical locations, can result from traumatic injury or disease. The most extreme manifestation of HO is represented by the rare, autosomal-dominant genetic disorder, fibrodysplasia ossificans progressiva (FOP), in which heterotopic bone forms progressively throughout the life of the individual, resulting in devastating effects on health, life expectancy and quality of life (*1, 2*). The great majority of FOP cases are caused by a single arginine to histidine amino acid substitution (R206H) in the intracellular glycine/serine-rich domain of the type 1 bone morphogenetic protein (BMP) receptor ACVR1 (*3, 4*). This amino acid substitution not only renders cells mildly hypersensitive to BMP ligands (*5, 6*) but also alters signaling specificity such that SMAD 1/5/8 phosphorylation and osteogenic differentiation is elicited by activin ligands (*7–9*), which normally activate SMAD 2/3 and function as inhibitors of BMP signaling (*7, 10, 11*). In fact, recent data has shown that activin A is both necessary and sufficient for HO flares, and its inhibition suppresses growth of nascent lesions in mouse models of FOP (*7, 9, 12, 13*). The safety and efficacy of systemic delivery of anti-activin A neutralizing antibodies are currently being evaluated in individuals with FOP (Clinical Trial # NCT03188666).

The relationship between physiological triggers of HO (see ref. (*14*), for review), such as soft tissue injury, inflammation, and immune cell activation, and inappropriate activation of ACVR1(R206H) by osteogenic ligands in cells responsible for heterotopic skeletogenesis, remains obscure. Also unexplained is the cause of so-called spontaneous HO, in which episodes of HO lesion formation (flare-ups), occur in the absence of known triggers (*14*). In this regard, a comprehensive natural history study indicates that spontaneous flare-ups account for at least half of all HO (*15*). To interrogate these and other disease pathophysiological mechanisms, and to test potential therapeutic agents and approaches, we developed a knockin mouse model of FOP (*Acvr1^tnR206H^*) in which the FOP-causing gene is expressed under the control of the endogenous *Acvr1* promoter upon Cre-dependent removal of the floxed stop cassette upstream of the mutant exon (*9*). Lineage-tracing studies in FOP mice support the conclusion that fibro/adipogenic progenitors (FAPs) – originally defined as muscle-resident bipotent mesenchymal progenitors (*16, 17*) – are a key offending cell type for both injury-induced and spontaneous disease (*9, 12, 18, 19*). Targeting *Acvr1^R206H^* expression to *Pdgfra*-positive cell populations, which in the postnatal musculoskeletal system are highly enriched for FAPs (*16, 20, 21*) and also includes periosteal cells (*9*), recapitulates HO at essentially all anatomical sites and tissues that affect individuals with FOP, including back and appendicular musculature, tendons and ligaments, major joints, and the jaw (*12*). Identification of skeletogenic progenitors responsible for HO now allows consideration of cell-specific therapies for FOP, such as gene therapies or other therapeutic interventions that alter the developmental capacity specifically of offending cell types.

In mice, as in humans, FOP mutations in *Acvr1* result in gain-of-function receptor activity that is manifested in the heterozygous state. Notably, in FOP mice, genetic loss of the remaining wild-type *Acvr1* allele in Tie2+ skeletogenic progenitors (predominantly FAPs) causes a profound increase in HO volume following muscle injury (*9*). These data suggest that the relative stoichiometry of wild-type and mutant ACVR1 receptors dictates disease severity and suggest a novel potential therapy for FOP based on over-expression of the wild-type ACVR1 receptor to dampen the osteogenic response, a possibility evaluated in the current study. To this end, we developed knockin mice that conditionally over-express a wild-type human *ACVR1* cDNA (*R26^nACVR1^*) and assessed whether *ACVR1* over-expression mitigates injury-induced and spontaneous HO in FOP mice. We also tested the capacity of global over-expression of *ACVR1* during prenatal development to rescue perinatal lethality of mice that express *Acvr1^R206H^* under endogenous transcriptional control (*22*). Collectively, the data demonstrate that over-expression of wild-type *ACVR1* is well tolerated and effectively dampens the pathological effects of the mutant FOP receptor.

## Results

### Characterization of mice that conditionally over-express human ACVR1

To produce mice that conditionally over-express the wild-type ACVR1 receptor, the human *ACVR1* cDNA was knocked into the constitutively expressed *Rosa26* locus (*23*), using a strategy and design similar to that used to produce the Cre-dependent eGFP reporter, *R26^NG^* (*24*). A floxed *Neo* cassette was positioned upstream of *ACVR1*, and the constitutively active CAG promoter/enhancer sequence (*25*) was provided to ensure high-level expression. Finally, an in-frame hemagglutinin sequence (HA) was added at the 3’ end of *ACVR1* to allow discrimination between human and mouse proteins, which are >98% identical. The unrecombined, *Neo*-containing, allele is designated *R26^nACVR1^*, whereas *R26^ACVR1^* denotes the Cre-recombined allele (Fig. S1A).

One *R26^nACVR1^* mouse line was established and maintained by routine breeding schemes. To validate expression of human *ACVR1* from the R26*^nACVR1^* allele, *R26^nACVR1^* mice were crossed with Tie2-Cre mice (*26*), and the HA peptide was detected by immunofluorescence on skeletal muscle sections derived from adult *Acvr1^tnR206H/+^*;*R26^nACVR1/+^*;Tie2-Cre mice. As expected, based on the known expression of *Tie2* (*21, 26, 27*), the HA tag was immunodetected in the vasculature and in a fraction of interstitial mononuclear cells [referred to as Tie2+ progenitors (*21*) or FAPs (*9*)], but not in muscle fibers, which strongly expressed tdTomato from the unrecombined *Acvr1^tnR206H^* allele (Fig. S1B, C). Quantification of *ACVR1* over-expression is provided below.

As BMP signaling regulates development of many tissues, we first determined whether widespread over-expression of *ACVR1* is compatible with viability. Males carrying the *R26^nACVR1^* allele were crossed with *Hprt^iCre/+^* (*28*) females to generate embryos in which the *R26^nACVR1^* allele was globally recombined (*R26^ACVR1^*). Although most *Hprt^iCre^* hemizygous male mice died before weaning irrespective of inheritance of *R26^nACVR1^*, *R26^ACVR1/+^*;*Hprt^iCre/+^* females developed to adulthood and were fertile. For subsequent analyses, both male and female *R26^ACVR1/+^* mice (germline-recombined at the *R26^nACVR1^* locus and lacking *Hprt^iCre^*) were generated using standard breeding schemes. The *R26^ACVR1^* allele was transmitted to offspring of weaning age at Mendelian frequencies (50.8%; 62 of 122 pups), and these *R26^ACVR1/+^* mice were fertile and did not show any gross physical or behavioral abnormalities.

*Acvr1*-null embryos exhibit multiple developmental defects and die during gastrulation (*29–31*). To test whether *R26^ACVR1^* can functionally replace the endogenous *Acvr1* gene, animals were produced that are homozygous null for *Acvr1* and heterozygous for *R26^ACVR1^*. Two null alleles were utilized: the unrecombined *Acvr1^tnR206H^* allele (*9*), and a second, independently-derived null allele (referred to here as *Acvr1^loxP^*) produced by crossing a floxed conditional *Acvr1* allele (*32*) (referred to here as *Acvr1^flox^*) with *Hprt^iCre^*. Notably, *Acvr1^loxP/loxP^*;*R26^ACVR1/+^*, *Acvr1^tnR206H/tnR206H^*;*R26^ACVR1/+^*, and *Acvr1^tnR2006H/loxP^*;*R26^ACVR1/+^* mice were viable and fertile and recovered at Mendelian frequencies at weaning (Table 1). These data demonstrate that the *R26^ACVR1^* allele can functionally replace the endogenous *Acvr1* gene.

**Table 1.** The *R26^ACVR1^* allele can functionally substitute for the endogenous *Acvr1* gene. Mice were generated by crossing *Acvr1^+/-^;R26^ACVR1/+^* and *Acvr1^+/-^* parents. Two Acvr1 null alleles were used (*Acvr1^tnR206H^* and *Acvr1^loxP^*) and are collectively designated as *Acvr1^-^*. Expected percentages are based strictly on predicted Mendelian frequencies, whereas Adjusted Expected percentages account for the known embryonic lethality of *Acvr1^-/-^* mice and assume complete rescue by *R26^ACVR1^*.

| *R26* genotype | *R26* wild type |  |  | *R26<sup>ACVR1</sup>* heterozygote |  |  |
| --- | --- | --- | --- | --- | --- | --- |
| *Acvr1* genotype | *Acvr1<sup>+/+</sup>* | *Acvr1<sup>+/-</sup>* | *Acvr1<sup>-/-</sup>* | *Acvr1<sup>+/+</sup>* | *Acvr1<sup>+/-</sup>* | *Acvr1<sup>-/-</sup>* |
| # mice at weaning | 3 | 17 | 0 | 5 | 10 | 7 |
| Expected (%) | 12.5 | 25 | 12.5 | 12.5 | 25 | 12.5 |
| Adjusted Expected (%) | 14.3 | 28.6 | 0 | 14.3 | 28.6 | 14.3 |
| Actual (%) | 7.1 | 40.5 | 0 | 11.9 | 23.8 | 16.7 |

### Over-expression of ACVR1 rescues perinatal lethality of FOP mice

Progeny that are heterozygous for the R206H-encoding germline mutation in *Acvr1* die perinatally (*19, 22*). To further characterize the effects of *Acvr1^R206H^* expression, the *Acvr1^tnR206H^* allele was globally recombined in the early embryo so that the pattern and timing of *Acvr1^R206H^* expression were dictated exclusively by *Acvr1* promoter activity. *Acvr1^tnR206H/+^* males were crossed with females that were heterozygous for the ubiquitously expressed *Hprt^Cre^* allele, and skeletal development and survival were assessed. Unlike the *Hprt^iCre^* allele used above, recombination of the paternally inherited *Acvr1^tnR206H^* allele after fertilization was not dependent on inheritance of the *Hprt^Cre^* allele, presumably because of accumulation of Cre in the egg (*33*). Thus, globally recombined offspring had the genotype of *Acvr1^R206H/+^*, *Acvr1^R206H/+^*;*Hprt^Cre/+^* (females), or *Acvr1^R206H/+^*;*Hprt^Cre^*/Y (males). Mice were born at the expected Mendelian frequency, but all pups carrying the recombined *Acvr1^R206H^* allele (17 of 34 neonates) died within 24 hours of birth and exhibited grossly visible malformation of the hindlimbs (Fig. 1F). Although the precise cause of death was not determined, affected pups lacked a milk spot in their bellies, raising the likelihood that death resulted from dehydration and nutritional deficits.

**Figure 1.**
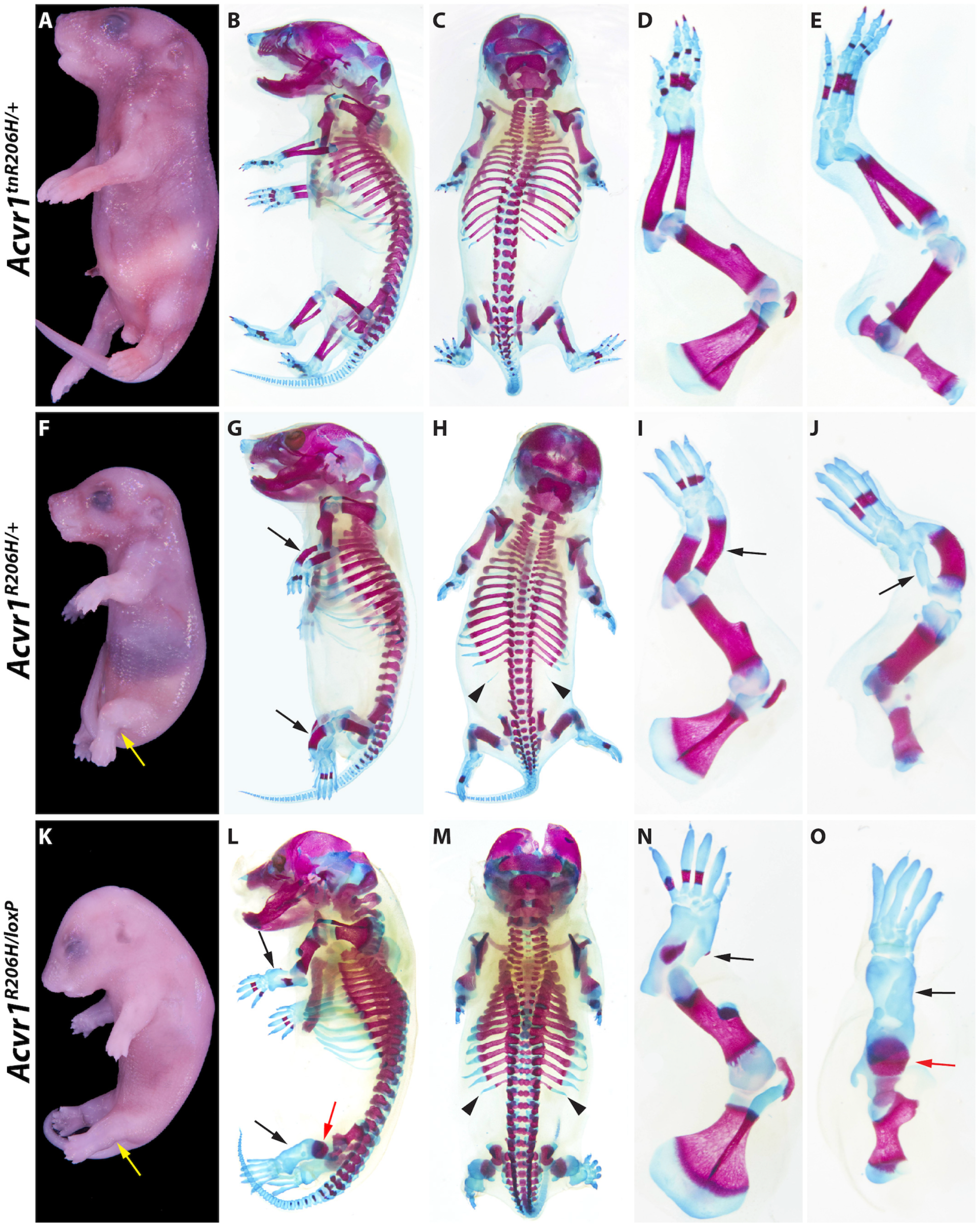
Representative examples of severe skeletal malformations caused by prenatal expression of *Acvr1^R206H^*. **(A-E)** *Acvr1^tnR206H/+^* control P0 neonate. (F-J) *Acvr1^R206H/+^* neonate. **(K-O)** *Acvr1^R206H/loxP^* neonate. Developmental defects are aggravated by loss of the wild-type *Acvr1* allele. **(A, F, K)** Whole-mount views after skin removal. Yellow arrows indicate abnormal positioning of the hindlimbs. Left side (B, G, L) and dorsal (C, H, M) views of ABAR-stained whole-mount neonates. ABAR-stained left forelimbs (D, I, N) and hindlimbs (E, J, O). Black and red arrows, severely shortened zeugopodal bones and femur, respectively; black arrowheads, L1 supernumerary ribs. Also note missing or abnormal ossification centers and thickened ribs in *Acvr1^tnR206H/+^* and *Acvr1^R206H/loxP^* neonates. Additional defects are described in the text.

Newborn *Acvr1^R206H/+^* mice did not exhibit HO, but Alcian blue/Alizarin red (ABAR) staining of P0 pups revealed pronounced skeletal defects, particularly of the forelimbs and hindlimbs (Fig. 1G-J) (n = 17). Bones of the hindlimb zeugopod (tibia/fibula) were grossly misshapen, the diaphyses were shortened and thickened, and the fibula lacked an ossification center (Fig. 1G-J). The digits of both the forelimb and hindlimb lacked distal ossification centers, the proximal ossification centers of digits 1 and 5 (forelimb) or digit 5 (hindlimb) were absent (Fig. 1I, J), most digital joints were absent, and cutaneous syndactyly was evident (Fig. 1F-J and Fig. S2A-D). Shortening of the fibula and defects in the formation of the metacarpophalangeal joints were already evident in embryos by 13.5 dpc (Fig. S2G, H). The axial skeleton also showed abnormalities, including rib thickenings in all *Acvr1^R206H/+^* P0 mice (Fig. 2G, H; Fig. S3C), the presence of supernumerary ribs associated with the first lumbar vertebrae (L1) in the majority of these mice, and spina bifida (Fig. 1H; Fig. S3C; n = 20 mice). Mice lacking the remaining wild-type *Acvr1* allele (*Acvr1^R206H/loxP^*) survived to birth, demonstrating that *Acvr1^R206H^* can fulfill the requirement of *Acvr1* in gastrulation (*29–31*). Importantly, however, loss of the wild-type *Acvr1* allele produced even more pronounced skeletal abnormalities and live newborns were not recovered. Skeletal defects included more severely truncated and malformed stylopods and zeugopods, further disruption of ossification including complete loss of autopod ossification in the hindlimbs, and thicker ribs (Fig. 1L-O). The exacerbated phenotype of *Acvr1^R206H/loxP^* mice suggests that wild-type ACVR1 partially suppresses the pathogenic developmental effects of ACVR1(R206H) in *Acvr1^R206H/+^* cells, a conclusion consistent with our previous findings (*9*). Perinatal lethality and most aforementioned skeletal malformations were fully penetrant, providing robust phenotypic criteria by which to evaluate the ability of *ACVR1* over-expression to mitigate the deleterious effects of pre-natal *Acvr1^R206H^* expression.

**Figure 2.**
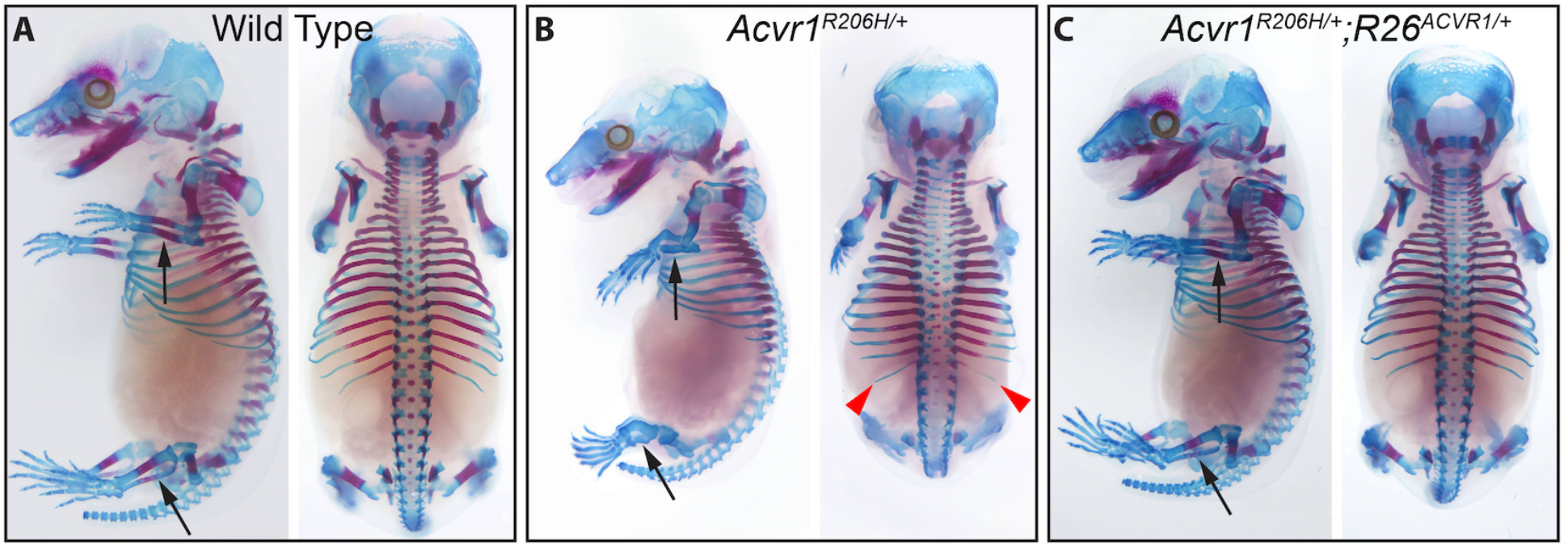
Suppression of *Acvr1^R206H^*-induced skeletal defects by over-expression of wild-type *ACVR1*. **(A-C)** Left and dorsal views of ABAR-stained 16.5 dpc fetuses. (A) Wild-type, (B) *Acvr1^R206H/+^* and (C) *Acvr1^R206H/+^*;*R26^ACVR1/+^* fetuses. Black arrows, zeugopod; red arrowheads, L1 supernumerary ribs.

To determine whether *ACVR1* over-expression can rescue lethality of FOP neonates, *Acvr1^tnR206H/+^*;*R26^nACVR1/+^* males were crossed to *Hprt^iCre/+^* females, and offspring were genotyped at weaning. As expected, no *Acvr1^R206H/+^* mice lacking *R26^ACVR1^* were recovered. Notably, however, mice carrying both the globally recombined *Acvr1^R206H^* and *R26^ACVR1^* alleles were viable and exhibited no externally visible abnormalities at weaning. Subsequent breeding of *Acvr1^R206H/+^*;*R26^ACVR1/+^* and wild-type mice confirmed *R26^ACVR1^*-dependent survival of *Acvr1^R206H/+^* mice, which were recovered at expected Mendelian frequencies at weaning (Table 2). Using several crossing schemes, a total of 90 adult mice were produced that were heterozygous for both the *Acvr1^R206H^* and *R26^ACVR1^* alleles. Except for one mouse that died prematurely (see below), mice showed no reduction in lifespan up to the latest experimental endpoint of 17-months-of-age, and mice were fertile. Skeletal development showed a concomitant normalization, as assessed by ABAR staining of 16.5 dpc fetuses (Fig.2A-C) and P0 pups (not shown). While supernumerary ribs at L1 were present in approximately 40% of cases (left and right sides scored separately; 16 of 27 mice), *ACVR1* over-expression rescued the rib thickening malformations and zeugopodal bone defects of the appendicular skeleton in all mice (Fig. 2C; Fig. S3D). Global expression of *R26^ACVR1^* also rescued the perinatal-lethal phenotype of the more severely affected *Acvr1^R206H/-^* mice (either *Acvr1^R206H/loxP^* or *Acvr1^R206H/tnR206H^*) (n=21 at weaning age or older; Fig. 3J).

**Figure 3.**
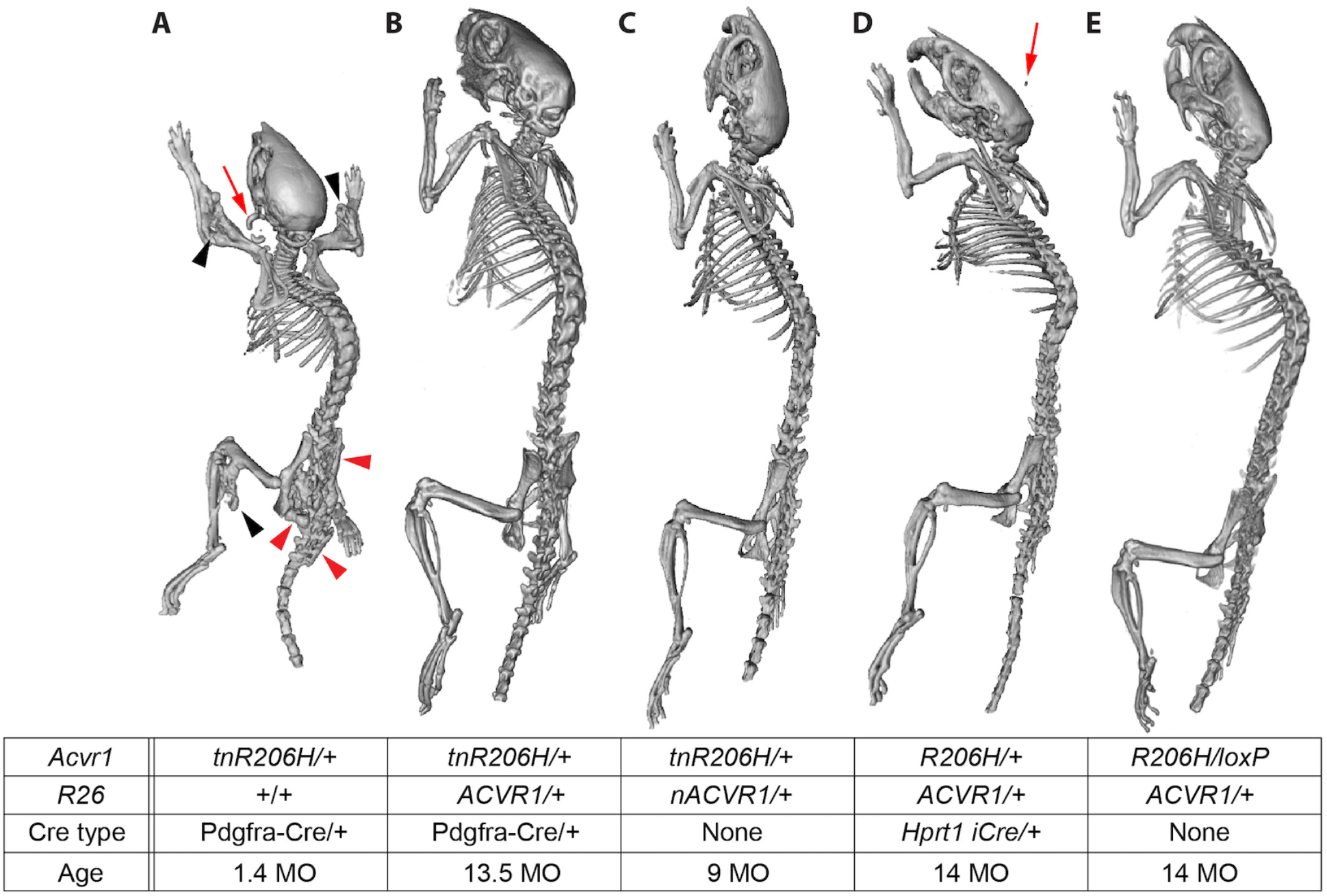
Suppression of spontaneous HO by over-expression of wild-type *ACVR1* in two models of FOP. The table shows the genotype and age (months) of each mouse. **(A-E)** Representative µCT images of juvenile (A) and adult (B-E) mice. Note that a juvenile *Acvr1^R206H/+^*;Pdgfra-Cre mouse (A) was used because of the poor survival and health of adult mice of this genotype. A control mouse that lacked Cre is shown in (C). The mouse in (E) inherited the recombined alleles shown through the germline, and their expression was therefore independent of Cre. Black and red arrowheads, spontaneous ectopic bone growth associated with the appendicular and axial skeleton, respectively; red arrows, HO at the ear-tag site.

**Table 2.** The *R26^ACVR1^* allele can rescue perinatal lethality of *Acvr1^R206H/+^* mice. Mice were generated by crossing *Acvr1^R206H/+^;R26^ACVR1/+^* and wild-type parental mice. Expected percentages are based strictly on predicted Mendelian frequencies, whereas Adjusted Expected percentages account for perinatal lethality of *Acvr1^R206H/+^* mice and assume complete rescue by *R26^ACVR1^*.

| Genotype | Wild type | *Acvr1<sup>R206H/+</sup>* | *R26<sup>ACVR1/+</sup>* | *Acvr1<sup>R206H/+</sup>;R26<sup>ACVR1/+</sup>* |
| --- | --- | --- | --- | --- |
| # mice at weaning | 22 | 0 | 19 | 19 |
| Expected (%) | 25.0 | 25.0 | 25.0 | 25.0 |
| Adjusted Expected (%) | 33.3 | 0.0 | 33.3 | 33.3 |
| Actual (%) | 36.7 | 0.0 | 31.7 | 31.7 |

### Inhibition of juvenile-onset spontaneous HO in FOP mice

Patient histories indicate that most new bone growth in FOP patients occurs spontaneously, without a known injury or inflammatory trigger (*15*). In previous analyses in which *Acvr1^tnR206H^* recombination was targeted to *Pdgfra*-positive progenitors, FOP mice exhibited aggressive, juvenile-onset spontaneous HO that was fully penetrant, characterized by HO at multiple anatomical locations, and survival to a median age of 39 days (*12*). Here, we addressed whether *ACVR1* over-expression reduces the occurrence and severity of spontaneous disease. In the present study, 20 of 23 *Acvr1^tnR206H/+^*;Pdgfra-Cre mice imaged by micro-computed tomography (μCT) between 33- and 90-days-of-age showed heterotopic bone growth (Fig. 3A). Affected mice typically exhibited widespread HO, with most mice having lesions at both axial and appendicular anatomical locations, as previously described (*12*). Strikingly, all *Acvr1^tnR206H/+^*;Pdgfra-Cre mice carrying the *R26^nACVR1^* allele survived to adulthood and appeared healthy, with no cases of premature death (89 mice; up to 15-months-old at the time of sacrifice). 12 mice between 5-weeks and 29-months-of-age were imaged by μCT. No skeletal malformations were detected in eight mice (Fig. 3B), whereas three mice exhibited a small exostosis-like expansion of the distal tibia (imaged at 13-, 44- and 64-weeks-of-age) (Fig. S4F).

### ACVR1 over-expression confers long-term protection from spontaneous HO in Acvr1^R206H^ germline mice

We next assessed the degree to which over-expression of *ACVR1* mitigates spontaneous HO in mice carrying the germline-recombined alleles, *Acvr1^R206H^* and *R26^ACVR1^*. Among the 90 adult mice produced that were heterozygous for both the *Acvr1^R206H^* and *R26^ACVR1^* alleles (7-weeks-old or older), one mouse was found dead at 52-days-of-age and imaging revealed fusion between zygomatic and mandibular bones, but no other sites of spontaneous bone growth. Eight mice between 7- and 9-months-of-age were scanned for HO by μCT. Strikingly, none exhibited spontaneous HO or other skeletal abnormalities. Three of these mice showed a small degree of heterotopic bone growth associated with the ear tag site, presumably resulting from minor injury caused by ear tagging. 17 additional mice in this cohort were imaged at 12-months-of-age or older. Among this older group, five mice showed abnormal growth of the distal tibia (Fig S4A-E), and six mice had minor HO at the ear tag site (Fig. S4C, D). Importantly, these overgrowths were dependent on the presence of the *Acvr1^R206H^* allele, although it is unclear whether these exostoses occurred because of greater expression of *Acvr1^R206H^* or reduced expression of the *R26^ACVR1^* allele in periosteal cells or other responsible cell types. Comparisons could not be made to *Acvr1^R206H/+^* mice, as these mice die at P0 (see above), although severe spontaneous HO at least as pronounced as that exhibited by *Acvr1^tnR206H/+^*;Pdgfra-Cre mice (Fig. 3A; also see ref. (*12*)) would be expected. Collectively, these data demonstrate that *ACVR1* over-expression confers long-term protection from spontaneous HO at all anatomical sites that typify disease manifestations in FOP mouse models and human patients.

### Over-expression of ACVR1 in mesenchymal progenitors protects FOP mice from injury-induced heterotopic ossification

Using a Tie2-Cre transgenic driver (*26*) to direct recombination of *Acvr1^tnR206H^* to Tie2+ mesenchymal cells, the majority of which have the marker profile of FAPs (*9, 21, 27*), consistently results in robust HO following pinch- or cardiotoxin-induced muscle injury (*9*). To test the consequence of *ACVR1* over-expression, adult *Acvr1^tnR206H/+^*;Tie2-Cre mice that either carry or lack the *R26^nACVR1^* over-expression allele were subjected to cardiotoxin-induced injury of the tibialis anterior (TA) muscle, and HO evaluated by μCT. 17 of 20 *Acvr1^tnR206H/+^*;Tie2-Cre FOP mice assessed at approximately 3 weeks post-injury (20-24 days) developed extraskeletal bone, and many of them (n=11) exhibited exostosis originating from the tibia, often manifested as localized thickening of the tibia near the middle of the tibial shaft (Fig. 4A; Fig. S5A). Notably, HO was undetectable among *Acvr1^tnR206H/+^*;*R26^nACVR1/+^*;Tie2-Cre mice and exostoses were absent (n=18) (Fig. 4B). Whole mount ABAR staining of 14 of these mice revealed that both heterotopic cartilage and bone were absent. We previously showed that genetic loss of the remaining wild-type *Acvr1* allele in *Acvr1^tnR206H^*^/*flox*^;Tie2-Cre mice causes profound exacerbation of injury-induced HO (*9*). ACVR1 over-expression also effectively blocked this extreme manifestation of trauma-induced HO (Fig. S5B, C).

**Figure 4.**
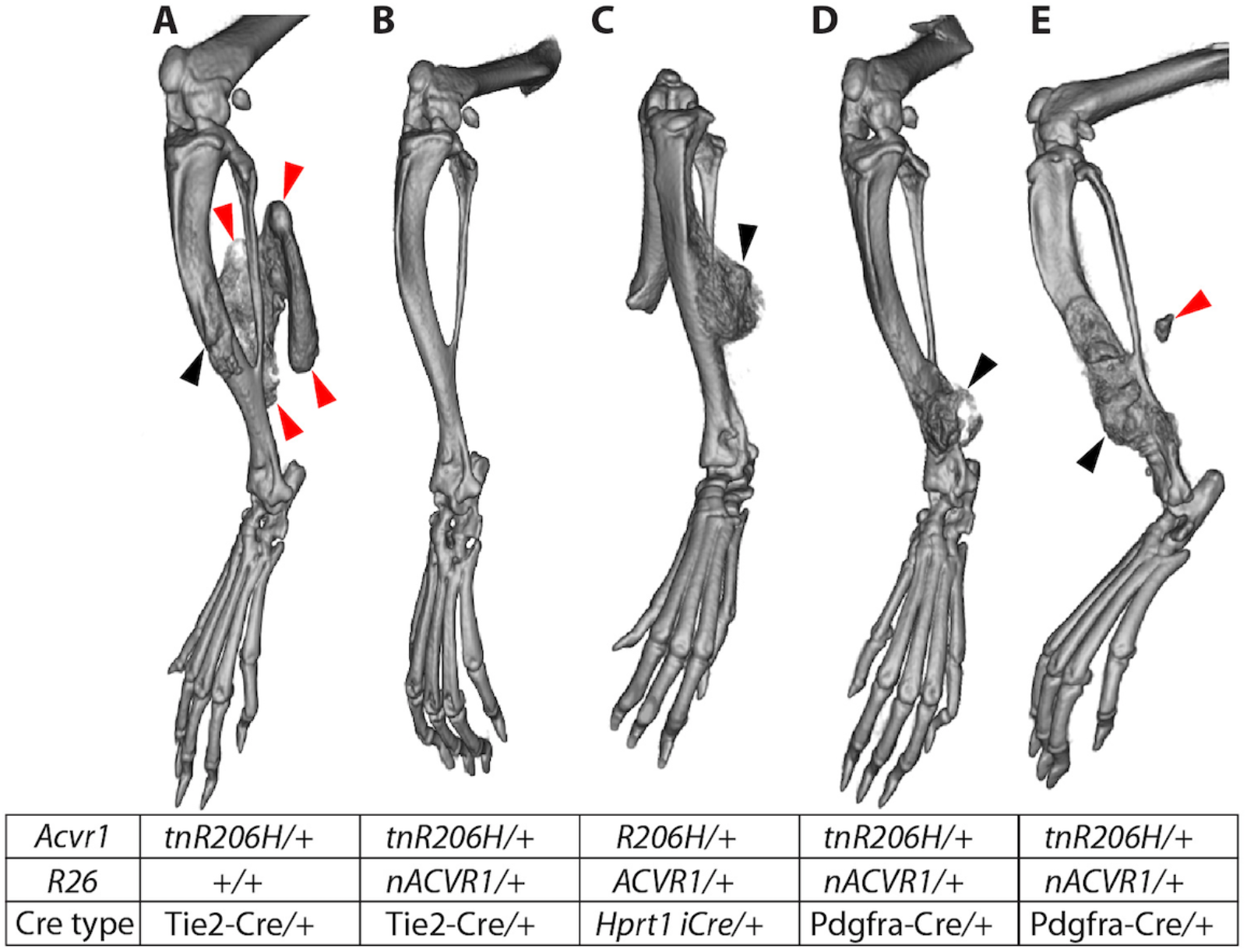
Inhibition of injury-induced HO by *ACVR1* over-expression. The table shows the genotype of each mouse. **(A-E)** µCT images captured 21-day after cardiotoxin injection of the TA muscle. Bone growth in (C) represents the most pronounced HO observed in mice of this genotype. Red arrowheads, HO that likely originated in soft tissue; black arrowheads, aberrant bone growth, apparently originating from the tibial surface. In one *Acvr1^tnR206H/+^*;*R26^nACVR1/+^*;Pdgfra-Cre mouse (E), a small volume of heterotopic bone was observed in the soft tissue near, but well-separated from, the tibia.

We typically do not use adult *Acvr1^tnR206H/+^*;Pdgfra-Cre mice as a model of injury-induced HO, both because aggressive spontaneous HO that begins by 4-weeks-of-age (*12*) complicates analyses, and because obtaining healthy adult mice in sufficient numbers is difficult due to their poor fitness and survival (*12*)(present study). Of note, however, only one of six adult *Acvr1^tnR206H/+^*;Pdgfra-Cre mice carrying the *R26^nACVR1^* allele developed extraskeletal bone after cardiotoxin-induced injury of the TA muscle (Fig. 4E), and this heterotopic bone was minor compared to the typical HO response of *Acvr1^tnR206H/+^*;Tie2-Cre mice (Fig. 4A). However, exostoses at the mid to distal tibia was observed in all *Acvr1^tnR206H/+^*;Pdgfra-Cre;*R26^nACVR1/+^* mice following injury of the TA muscle (Fig. 4D, E; n=6). Remarkably, even germline-recombined *Acvr1^R206H/+^*;*R26^ACVR1/+^* mice did not develop extraskeletal bone after cardiotoxin-induced injury of the TA muscle (n=10). The only evident abnormality, which ranged in severity but was often pronounced, was localized outgrowth of endogenous bone in most mice (7 of 10 mice), sometimes extending from the tibia and invading soft tissues (Fig. 4C).

### Quantification of mRNA from the R26^ACVR1^ and endogenous Acvr1 alleles

The functionality of the *R26^ACVR1^* allele was shown by its ability to substitute for the endogenous *Acvr1* gene and to rescue perinatal lethality of *Acvr1^R206H/+^* mice. To quantify expression from the *R26^ACVR1^* allele, mRNA was isolated from adherent unfractionated cells derived from hindlimb muscles and associated soft tissues of adult *Acvr1^R206H/+^*;*R26^ACVR1/+^* mice (these include FAPs, as well as other uncharacterized mesenchymal cell types; unpublished observations), and mRNA quantified by RT-droplet digital PCR (ddPCR) with primers that distinguish transcripts from the *Acvr1^R206H^*, wild-type *Acvr1*, and *R26^ACVR1^* alleles. Human *ACVR1* mRNA was 40- to 50-fold more abundant than endogenous *Acvr1^R206H^* and *Acvr1* mRNAs, which were equivalently expressed (Fig. 5A), and abundance of the endogenous alleles was not affected by the presence of *R26^ACVR1^* (Fig. 5A, B). A similar ratio of exogenous to endogenous transcripts was observed in 14.5 dpc *Acvr1^R206H/+^*;*R26^ACVR1/+^* mouse fetuses (Fig. 5C).

**Figure 5.**
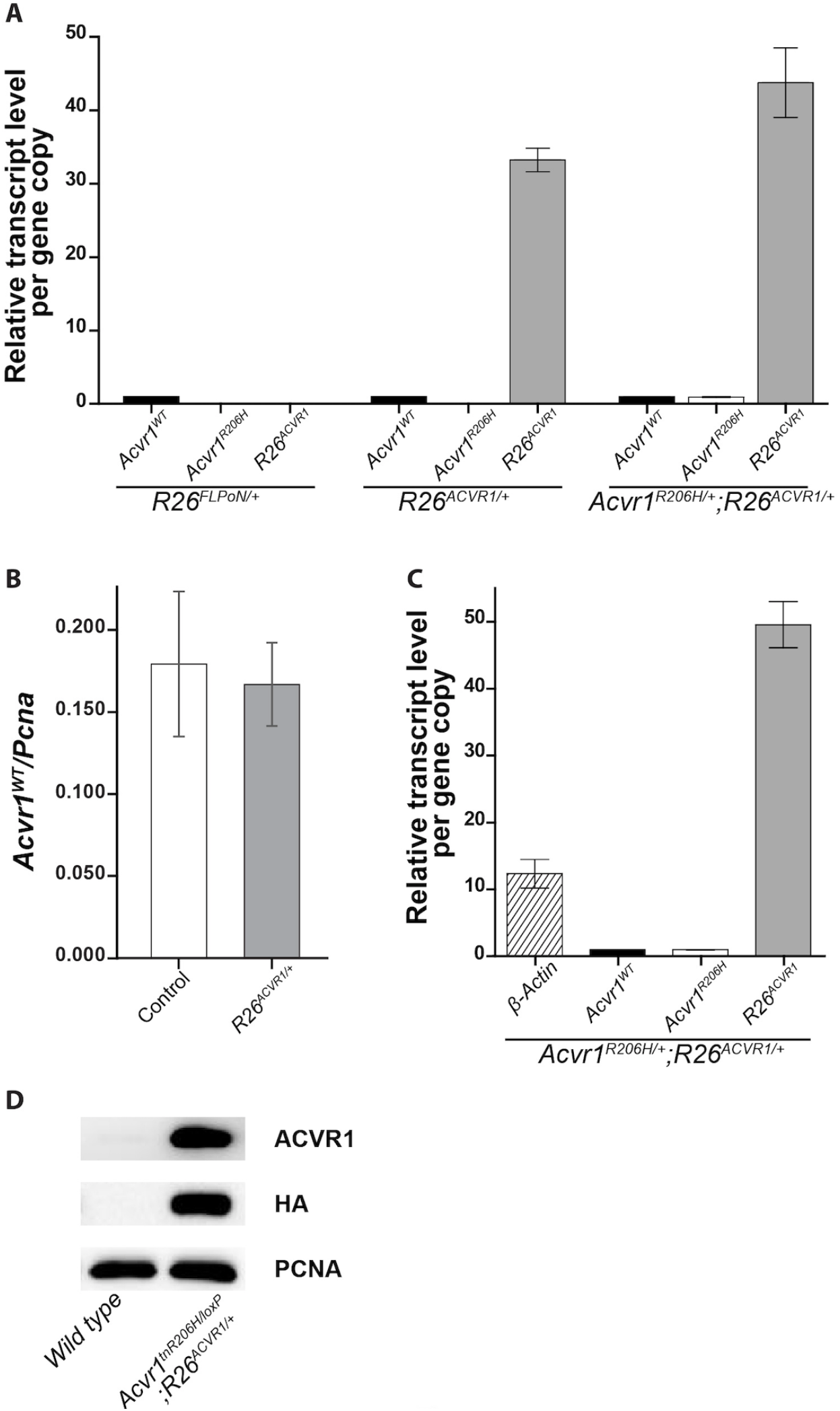
Quantification of exogenous ACVR1 mRNA and protein. **(A, C)** RT-ddPCR quantification of *ACVR1* mRNA from cultured adherent cells derived from total adult leg muscles and surrounding soft tissues (A), or whole 14.5 dpc fetuses (C). In (A), *R26^FLPo/+^* mice served as a control for possible non-specific effects of a gene knockin at the *Rosa* locus. The transcript level of each gene allele was normalized to that of *Acvr1^WT^*. **(B)** Effect of *R26^ACVR1^* on transcriptional activity of the endogenous *Acvr1* gene. The transcript level of *Acvr1^WT^* was normalized to *Pcna*, which was used as an internal control. **(D)** Western blot of HA-tagged ACVR1 protein from lysates of cultured adherent cells derived from total adult leg muscles and surrounding soft tissues. Antibody specificity is shown in the right column. Note that *Acvr1^tnR206H/loxP^* cells cannot produce functional full-length ACVR1 protein; the signal detected with the anti-ACVR1 antibody represents exogenous ACVR1 only. Endogenous ACVR1 in wild-type cells is barely detectable under these conditions.

ACVR1 protein showed a strong signal on Western blots of extracts from *R26^ACVR1^*-expressing adherent cells using an anti-ACVR1 antibody (Fig. 5D). As human and mouse ACVR1 proteins could not be distinguished with anti-ACVR1 antibodies, analysis of exogenous ACVR1 expression utilized cells lacking a functional *Acvr1* allele (derived from *Acvr1^tnR206H/loxP^*;*R26^ACVR1/+^* mice). Whereas the signal with an anti-ACVR1 antibody reflected a marked increase in protein abundance in *R26^ACVR1^*-expressing cells compared to wild-type cells, the calculated increase varied substantially between biological replicates (17-fold and 57-fold), likely because the ACVR1 signal in wild-type cells was barely detectable (Fig. 5D). Expression of exogenous ACVR1 was confirmed using an anti-HA antibody (Fig. 5D), which detects the ACVR1-HA fusion protein.

### Effects of ACVR1 over-expression on activin A-induced osteogenic signaling

We next tested whether over-expression of wild-type *ACVR1* affects pSMAD1/5/8 osteogenic signaling of *Acvr1^R206H^*-expressing cells in response to activin A and BMP6 ligands. Whereas BMP6 can signal through ACVR1 and activate osteogenic signaling of both wild-type and FOP cells, most studies have shown that osteogenic signaling in response to activin A is dependent on ACVR1(R206H) (*7–11*). Fluorescence-activated cell sorting was used to isolate FAPs from total hindlimb muscles of control mice (*R26^NG/+^*), *ACVR1* over-expressing mice (*R26^ACVR1/NG^*), and FOP mice that either carried (*Acvr1^tnR206H/+^*;*R26^ACVR1/NG^*;Tie2-Cre) or lacked (*Acvr1^tnR206H/+^*;*R26^NG/+^*;Tie2-Cre) the germline transmitted *R26^ACVR1^* allele, as previously described (*9, 21, 27*). Cultured FAPs were serum-starved for 2 hrs, treated with different concentrations of activin A or BMP6 for 1 hr, and pSMAD1/5/8 quantified by Western blotting. Consistent with previous results, R206H-FAPs, but not control FAPs lacking *Acvr1^tnR206H^*, induced SMAD1/5/8 phosphorylation in response to activin A (*9*), and the response was dose-dependent (Fig. 6A, C). Importantly, *ACVR1* over-expression dampened the response to activin A, although the reduction in pSMAD1/5/8 levels was modest at the highest concentration of activin A (4 nM; 100 ng/mL) (Fig. 6A, C). As expected, BMP6 stimulated concentration-dependent SMAD1/5/8 phosphorylation of control FAPs. (Fig. 6B, D). Interestingly, however, over-expression of *ACVR1* did not sensitize cells to BMP6 at any ligand input concentration; in fact, pSMAD1/5/8 levels were modestly reduced in *ACVR1* over-expressing cells (Fig. 6B, D). The finding that *ACVR1* over-expression does not confer hyper-responsiveness to BMP6 suggests the limited availability of one or more positively-acting signaling components, such as obligatory type II receptors, or positively acting effectors downstream of receptor engagement.

**Figure 6.**
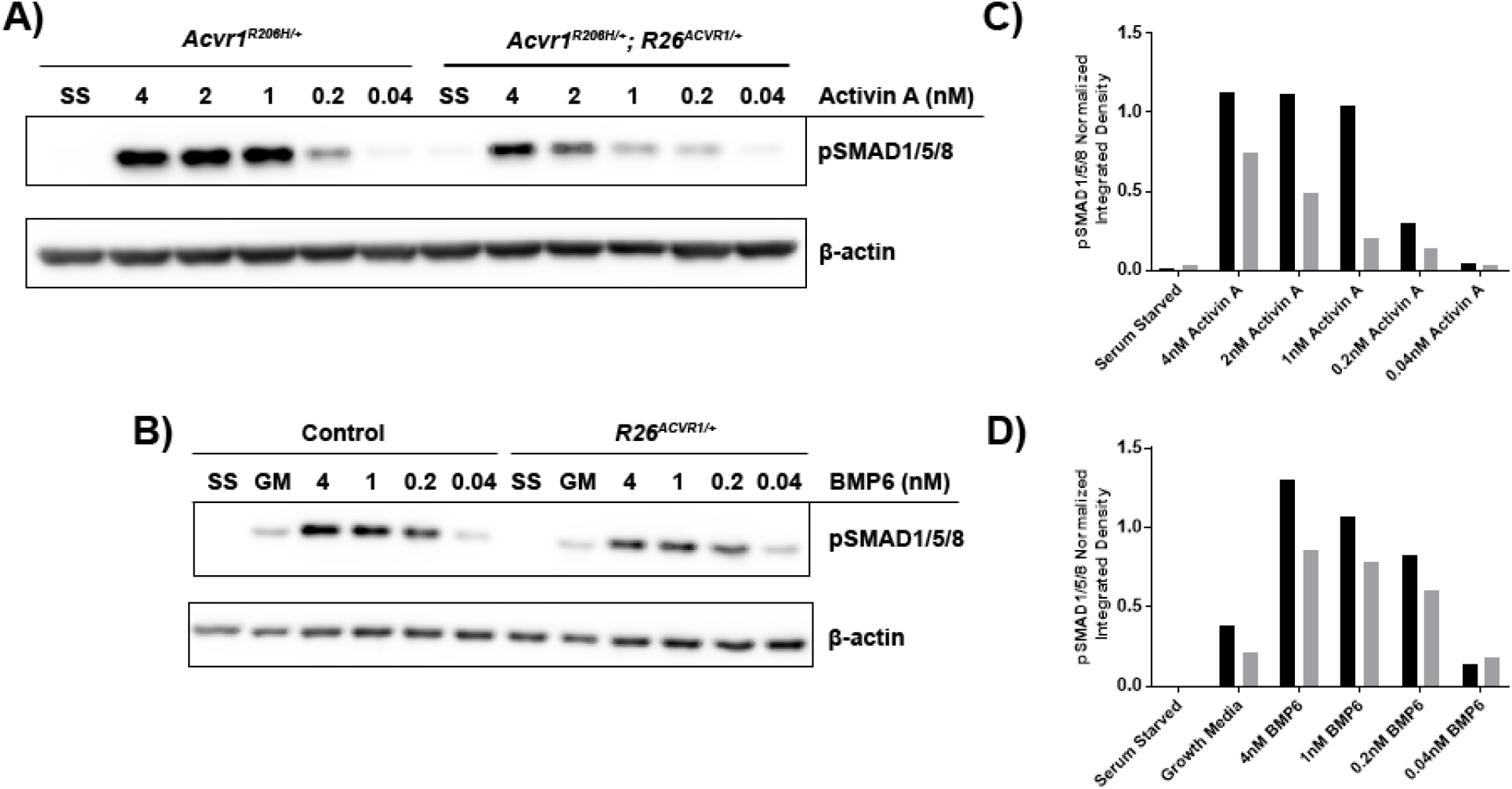
Effects of *ACVR1* over-expression on activin A- and BMP6-induced SMAD1/5/8 phosphorylation. **(A-D)** Representative western blots and corresponding quantification of pSMAD1/5/8 from lysates of FAPs of the indicated genotypes. After serum starvation for 2 hours, cells were incubated with the indicated concentrations of activin A (A) or BMP6 (B) for 1 hour prior to cell lysis. β-actin was used as an internal control to normalize pSMAD1/5/8 signal intensity. **(C)** pSMAD1/5/8 quantification of the blot in (A). **(D)** pSMAD1/5/8 quantification of the blot in (B). In (C) and (D), the integrated densities of pSMAD1/5/8 and β-actin bands were quantified using ImageJ, and normalized pSMAD1/5/8 values are shown.

### ACVR1 over-expression reduces fibroproliferative cell expansion and BMP signaling following muscle injury

We next addressed whether inhibition of injury-induced HO by *ACVR1* over-expression was associated with reduced osteogenic signaling. For this analysis, the FOP mouse model *Acvr1^tnR206H/+^*;*R26^NG/+^*;Pdgfra-Cre was used, as the osteogenic response is typically greater than when recombination is driven by Tie2-Cre (*9, 12, 21*). Juvenile mice were used for this analysis to avoid the aforementioned complications of using adult *Acvr1^tnR206H/+^*;*R26^NG/+^*;Pdgfra-Cre mice for injury studies. First, the distribution of pSMAD1/5/8-positive cells was determined 3 days after injection of cardiotoxin into the TA muscle of FOP mice. With the relatively large volume of cardiotoxin used (100 μL), surrounding muscles, including the EDL, peroneus longus, plantaris and gastrocnemius, often showed varying extents of damage (unpublished observations). Interestingly, unlike induction of HO within the TA by intramuscular injection of BMP2 (*21*), HO typically developed posterior to, but separated from, the fibula, and on the surface of the tibia, rather than within the TA itself. This was also observed when the Tie2-Cre driver was used (Fig. 4A; Fig. S5A, B) and in a pinch-injury model (data not shown). As the ultimate anatomical location of HO varied between mice and likely emanated from connective tissues between muscles rather than intramuscularly, surveys of cryosections of the entire lower hindlimb at day 3 post-injury were undertaken to identify areas of mesenchymal hypercellularity. Irrespective of genotype (control, FOP, and *ACVR1* over-expressing mice), muscle injury resulted in increased mesenchymal cellularity (fibroproliferative response) of intermuscular fascia (typically, the posterior and transverse intermuscular septa) and loss of defined septal boundaries (Fig. 7A, B, D, E; Fig. S6). Importantly, however, FOP mice exhibited a much more pronounced fibroproliferative response, which is characteristic of early, pre-skeletal FOP lesions in mice and humans (*22, 34*) (Fig. 7A; Fig. 8A; Fig. S6B). Cells that stained intensely for both pSMAD1/5/8 and the cartilage/pre-cartilage marker SOX9, which is first detected prior to histologically identifiable cartilage (day 5 to 6 in this model), were abundant in the fibroproliferative regions of FOP mice (Fig. 7A, D; Fig. 8B, C). The majority of SOX9-positive cells expressed the *R26^NG^* lineage tracer and were negative for tdTomato (recombined at the *Acvr1^tnR206H^* locus), although some tdTomato-positive cells also stained for SOX9 (Fig. 8B-E). Whether these unrecombined SOX9-positive cells would have contributed to definitive HO at endpoint could not be determined, although our previous data indicated that the great majority of definitive HO cells express *Acvr1^R206H^* (*9*). Some mesenchymal cells within regenerating muscles located near fibroproliferative areas (typically, gastrocnemius and plantaris muscles) also expressed pSMAD1/5/8 and SOX9, although apparent protein levels were substantially lower than in mesenchymal cells of presumptive HO regions (Fig. 7G; Fig. 8F, G; Fig. S7G-I). In both control and FOP mice, centrally localized nuclei within nascent regenerating fibers stained weakly for pSMAD1/5/8 (compare Fig. 7B, C and Fig. S7A-C) and SOX9 (data not shown). SOX9 expression in wild-type muscle at this stage is consistent with a previous report of transient SOX9 expression during early myogenic differentiation (*35*).

**Figure 7.**
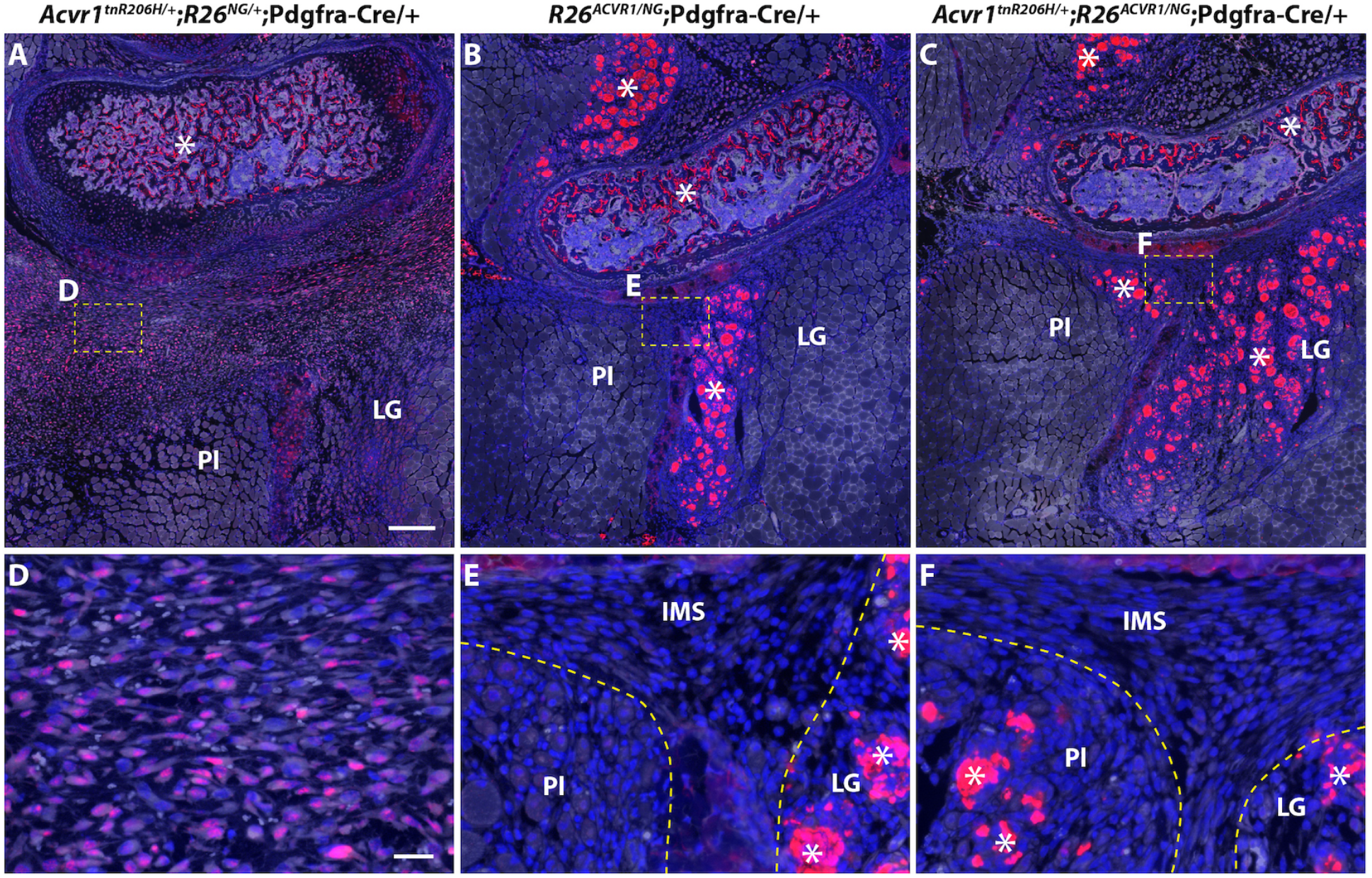
*ACVR1* over-expression greatly reduces the number of pSMAD1/5/8-positive cells in the hindlimbs of juvenile (33-day-old) mice 3 days after cardiotoxin-induced injury of the TA muscle. Fibroproliferation was observed in the intramuscular septa posterior to the fibula in all genotypes, but was most pronounced in *Acvr1^tnR206H/+^*;*R26^NG/+^*;Pdgfra-Cre mice. **(A-C)** Representative low magnification images of cross sections of the lower leg showing the presumptive HO-forming region located posterior to the fibula of a *Acvr1^tnR206H/+^*;*R26^NG/+^*;Pdgfra-Cre mouse, and the corresponding regions of *R26^ACVR1/NG^*;Pdgfra-Cre (B) and *Acvr1^tnR206H/+^*;*R26^ACVR1/NG^*;Pdgfra-Cre (C) mice. Boxed areas D, E, and F are shown at higher magnification in panels (D), (E), and (F). **(D-F)** Magnified views of fibroproliferative regions in the intermuscular septa corresponding to the boxed areas in (A-C). Note that GFP signal from the recombined *R26^NG^* reporter was destroyed during antigen retrieval for pSMAD1/5/8 detection. Cells with nuclear-localized pSMAD1/5/8 signal (red) were abundant in the intermuscular septum of *Acvr1^tnR206H/+^*; *R26^NG/+^*;Pdgfra-Cre mice (A, D), but not in *R26^ACVR1/NG^*;Pdgfra-Cre (B, E) or *Acvr1^tnR206H/+^*;*R26^ACVR1/NG^*;Pdgfra-Cre (C, F) mice. Autofluorescence through the green channel is shown in white to better reveal tissue structure. Damaged muscles are outlined by hatched lines (E, F). The strong non-nuclear fluorescent signal, which was present in all genotypes and considered non-specific staining, was observed in, but not limited to, damaged myofibers and the trabecular bone (examples at asterisks). Sections were counterstained with DAPI (blue).

**Figure 8.**
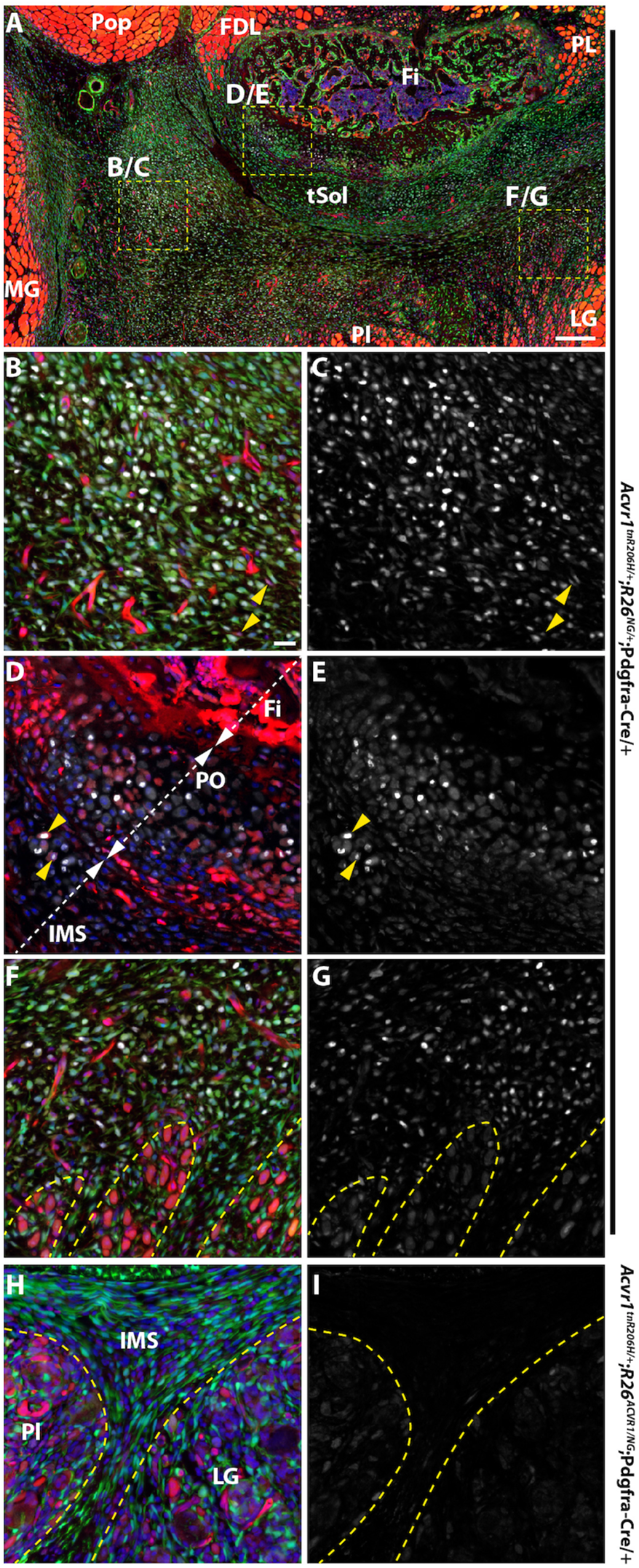
*ACVR1* over-expression inhibits induction of SOX9 in fibroproliferative cells in the hindlimbs of juvenile (33-day-old) mice after cardiotoxin-induced injury of the TA muscle. Tissue was collected 3 days post-injury. Markers are as follows: SOX9 (white, A-I); tdTomato from the unrecombined *Acvr1^tnR206H^* allele (red, A, B, D, F, H); GFP from the recombined *R26^NG^* reporter (green, A, B, F, H); DAPI (blue, A, B, D, F, H). **(A-G)** Representative images from an *Acvr1^tnR206H/+^*;*R26^NG/+^*;Pdgfra-Cre mouse. **(A)** Low magnification image of a cross section of the lower leg showing the presumptive HO-forming region located posterior to the fibula. Boxed areas correspond to regions shown at higher magnification in panels B/C, D/E, and F/G. **(B, C)** Portion of the fibroproliferative region of the transverse intermuscular septum. SOX9-positive cells are abundant. The great majority of SOX9+ cells are recombined at the *Acvr1^tnR206H^* locus, although a few unrecombined, tdTomato-positive cells are weakly positive for SOX9 (yellow arrowheads). Most of the red fluorescence in (B) represents unrecombined vascular elements. **(D, E)** High magnification view of a portion of the fibular periosteum (PO) and juxtaposed intermuscular septum (IMS). A small portion of the posterior fibula (Fi) is shown. Some cells of the periosteum are SOX9+. A small number of *Acvr1^tnR206H^*-unrecombined cells are positive for SOX9 (examples at yellow arrowheads). **(F, G)** Image representing the boundary between the SOX9+ fibroproliferative region and the regenerating lateral gastrocnemius. **(H, I)** Representative image from an *Acvr1^tnR206H/+^*;*R26^ACVR1/NG^*;Pdgfra-Cre mouse. The intermuscular septum is largely devoid of SOX9+ cells. Damaged muscles are outlined by hatched lines in (F-I). FDL, flexor digitorum longus; Fi, fibula; LG, lateral head of gastrocnemius; MG, medial head of gastrocnemius; Pl, plantaris; Pop, popliteus; tSol, soleus tendon; IMS, intermuscular septum. Images in (B-I) are at the same magnification.

FOP mice that over-express *ACVR1* also exhibited increased cellularity of intermuscular septa after injury of the TA muscle (Fig. 7C, F; Fig. 8H). Importantly, however, they lacked the pronounced fibroproliferative cell accumulations and strong pSMAD1/5/8 and SOX9 staining that typify early lesions in FOP mice (Fig. 7C, F; Fig. 8H, I). Like FOP and control mice, FOP mice carrying the *R26^ACVR1^* allele exhibited weak pSMAD1/5/8 and SOX9 staining of nascent myonuclei and intramuscular interstitial cells (Fig. S7C, F, I; data not shown). Collectively, these data show that *ACVR1* over-expression greatly diminishes osteogenic signaling and mesenchyme expansion in anatomical locations typically associated with HO, but does not affect BMP signaling in nascent muscle fibers in regenerating muscle.

## Discussion

The impetus to explore the possible therapeutic applicability of manipulating the stoichiometric balance of wild-type and mutant ACVR1 receptors was based on our previous finding that loss of the wild-type ACVR1 receptor in FOP mice profoundly exacerbated trauma-induced HO, which remained activin A dependent (*9*). Here, we developed knockin mice that conditionally over-express wild-type *ACVR1* and assessed whether over-production of the wild-type receptor in FOP mice has the opposite effect, namely, to mitigate the pathogenic effects of ACVR1(R206H). We showed that even global over-expression of human *ACVR1* is well-tolerated and has no obvious negative effects on development, longevity, or reproduction. This strategy was highly effective at mitigating or abrogating most of the deleterious effects of *Acvr1^R206H^* expression, including injury-induced and spontaneous HO, and the major developmental skeletal abnormalities and perinatal lethality of *Acvr1^R206H/+^* mice. Additionally, the protective effects of the *R26^ACVR1^* allele allowed us to generate the first adult mice that carry a germline *Acvr1^R206H^* mutation (see ref. (*22*)), providing the most rigorous and physiologically relevant demonstration of the therapeutic efficacy of *ACVR1* over-expression to date.

We note that while HO of apparent soft tissue origin was effectively inhibited in all FOP models, not all manifestations of abnormal bone growth were eliminated by *ACVR1* over-expression. In particular, exostosis associated with the tibial surface, which typically resulted in localized thickening of the tibia, was common following cardiotoxin-induced injury of the TA muscle of *Acvr1^tnR206H/+^*;*R26^nACVR1/+^*;Pdgfra-Cre and *Acvr1^R206H/+^*;*R26^ACVR1/+^* mice. In some *Acvr1^R206H/+^*;*R26^ACVR1/+^* mice, substantial ectopic bone growth apparently emanating from the tibial surface invaded the surrounding soft tissues. We cannot evaluate the degree to which *ACVR1* over-expression mitigated HO in these models because of the lack of comparison groups due to inviability (*Acvr1^R206H/+^*), or the complicating effect of aggressive, juvenile-onset spontaneous HO [*Acvr1^tnR206H/+^*;Pdgfra-Cre; (*12*) and present study]. We note, however, that the *R26^ACVR1^* allele did effectively suppress the tibial overgrowth phenotype in *Acvr1^tnR206H/+^*;Tie-Cre mice, even when this phenotype was greatly exacerbated by loss of the wild-type *Acvr1* allele. These data demonstrate that the tibial overgrowth phenotype can be blocked by *ACVR1* over-expression in certain experimental settings. As the specificity of Tie2-Cre and Pdgfra-Cre expression is overlapping but distinct (*9, 12, 21, 26, 36*), differences in *ACVR1* and/or *Acvr1^R206H^* expression levels in distinct cell types targeted by these Cre drivers is one possible explanation for apparent differences in efficacy of *ACVR1* over-expression in these models.

*ACVR1* over-expression probably dampens activin-dependent BMP signaling by reducing the absolute number or density of ligand-engaged ACVR1(R206H)-containing signaling complexes at the surface of skeletogenic progenitors. Activin A and other members of the TGFβ ligand superfamily signal through heterotetrameric complexes that are typically comprised of two type I and two type II receptor serine-threonine kinases. Members of the activin/inhibin and BMP/GDF sub-groups share type II receptors (ACVR2A, ACVR2B, BMPR2 for BMP/GDF ligands and predominately ACVR2A and ACVR2B for activin/inhibin ligands), and association with these type II receptors is required for type I receptor signaling activity (*37, 38*), including that of both ACVR1 and ACVR1(R206H)(*39–41*). In this context, competition between over-expressed wild-type ACVR1 and ACVR1(R206H) for limiting quantities of one or more obligate type II receptors is an attractive possibility, as it would directly result in fewer ACVR1(R206H)-containing tetrameric signaling complexes at the cell surface. Receptor complexes comprised of wild-type ACVR1 should serve as an activin A sink, since activin A binding stimulates very weak (*41, 42*) or no detectable (*7–11*) SMAD1/5/8 phosphorylation. Thus, if activin A is present at sub-saturating levels, the increased representation of wild-type ACVR1 should further reduce pathogenic signaling. Although the molecular details differ, a conceptually related phenomenon was recently described for the type I TGFβ receptor TGFβRI, which competes with the type I receptor ACVRL1 (ALK1) for ACTRIIB (ACVR2B) binding, thereby constraining BMP9 signaling in the growth plate through highly responsive ACVRL1/ACTRIIB complexes (*43*). More relatedly, competition for the shared type II receptor BMPRII probably explains the inhibitory effects of transfected wild-type ACVR1 on BMP2- and BMP4-dependent osteogenic signaling through BMPR1A and BMPR1B in cell culture models (*6*).

While many studies have demonstrated that BMP signaling is dysregulated in ACVR1(R206H)-expressing cells, less clear is the composition of receptor complexes that activate BMP signaling pathways in these cells. Given the promiscuity that characterizes interactions within and between the type I and type II receptor families (*38, 44, 45*), many distinct ACVR1(R206H)-containing complexes probably exist on the surface of heterozygous FOP cells. In terms of the type I receptor component, these include ACVR1(R206H) homodimers, and heteromers in which the type I partner is either wild-type ACVR1 or other type I receptors. Although definitive data is lacking, several observations point to receptor complexes containing ACVR1(R206H) homodimers as key drivers of HO in FOP. First, the greatly exacerbated HO phenotype of FOP mice lacking the wild-type *Acvr1* allele in skeletogenic progenitors (*9*) suggests that complexes comprised of ACVR1(R206H) homodimers exhibit increased or dysregulated BMP signaling activity compared to their ACVR1/ACVR1(R206H) heteromeric counterparts. Second, ACVR1(R206H) does not require typical type I receptor partners to activate pSMAD1/5/8. In zebrafish embryos, for example, Acvr1/Bmpr1 heteromeric complexes are required to transduce BMP signals in dorsoventral patterning (*46, 47*), whereas Bmpr1 is not required for signaling via exogenously provided ACVR1(R206H) in this model (*48*). Similarly, ACVR1B/C is required to induce SMAD1/5 phosphorylation by activin A in wild-type HEK293T cells, but these type I receptors are dispensable in *Acvr1^R206H/R206H^* cells (*41*). Third, although not demonstrated to be solely attributable to ACVR1(R206H) homomeric complexes, *Acvr1^R206H/R206H^* HEK293T cells exhibited a modestly extended duration of receptor activation in response to activin A (*41*). Finally, in contradistinction to wild-type cells, pSMAD1/5 activation driven by activin A-dependent receptor clustering did not require ACVR2A/B kinase activity in *Acvr1^R206H/R206H^* cells (*41*). Regarding the present study, probabilistic estimates predict that over-expression of wild-type *ACVR1* in FOP mice would most dramatically reduce representation of ACVR1(R206H) homodimeric complexes, under the reasonable assumption that wild-type and mutant ACVR1 receptors have similar binding affinities for receptor partners. From these observations, we hypothesize that tetrameric complexes containing ACVR1(R206H) homodimers represent the main drivers of pathological bone formation in FOP. By extension, it is reasonable to propose that *ACVR1* over-expression inhibits HO in FOP mice both by competing for essential signaling components and by titrating ACVR1(R206H) into inactive or less active heteromeric receptor complexes.

Given the essential roles of the TGFβ superfamily in mesoderm formation, skeletogenesis, and many other developmental and physiological processes (*49, 50*), one might predict that over-expression of *ACVR1* at levels reported here would have deleterious effects, which could result from any of several gain-of-function or dominant-negative mechanisms. Examples include sensitizing cells to BMP ligands that signal through ACVR1 (*37, 51*), reducing SMAD2/3 signaling by sequestration of activins into inactive complexes (*7, 11*), or dampening signaling by other TGFβ family ligands and type I receptors by reducing availability of shared type I or type II receptors (see ref. (*6, 37, 38, 46, 47*). Interestingly, however, over-expression of the *Acvr1* orthologs in the zebrafish early embryo (*52, 53*) or chicken limb bud (*54*) did not perturb development, or limb skeletogenesis, respectively. Consistent with these observations, we showed that global over-expression of human *ACVR1* rescued embryonic lethality of *Acvr1*-null mouse embryos, which normally die during gastrulation (*29–31*), and was well tolerated, having no obvious negative developmental effects. In addition, adult mice that globally over-express *ACVR1* did not develop HO in response to muscle injury. This lack of responsiveness was not due to competition between activin A and BMPs for binding to ACVR1-containing receptor complexes (see refs. (*7, 10, 11*)), as globally recombined *R26^ACVR1/+^* mice were refractile to injury-induced HO even under conditions of antibody-mediated inhibition of activin A (unpublished observations). These data could indicate that BMPs that signal through ACVR1, such as BMP5, BMP6 and BMP7 (*37, 55, 56*), are not available at sufficient levels in skeletal muscles and associated connective tissues to elicit an osteogenic response from presumptively hyper-responsive cells. Alternatively, *ACVR1* over-expression may not sensitize skeletogenic progenitors to BMP ligands, perhaps because of insufficient quantities or activities of one or more BMP signal transduction components. The latter possibility is consistent with western blot analyses, which showed that *ACVR1* over-expressing FAPs did not exhibit enhanced SMAD1/5/8 phosphorylation at any BMP6 input concentration tested. To the contrary, pSMAD1/5/8 levels were modestly reduced in *ACVR1* over-expressing cells.

To date, three pharmacological treatment modalities have shown promise in pre-clinical models of FOP, all of which involve systemic inhibition of signaling pathways involved in diverse cellular processes and tissue homeostasis (*57*). These include anti-activin A antibodies to block productive ligand-receptor interactions, small molecule inhibitors of ACVR1 kinase activity, and inhibition of cartilage differentiation by retinoic acid receptor gamma (RARγ) agonists (*37, 57*). Recently, a clinical trial with the RARγ agonist palovarotene (Ipsen: NCT03312634) was placed on hold for children under the age of 14 because of incidents of premature growth plate closure, a consequence consistent with the known toxic effects of palovarotene in FOP mice (*12, 58*) (but see ref. (*59*)). Antibody-mediated inhibition of activin A has shown promising results in a Phase II clinical trial for adult patients with FOP (Regeneron: NCT03188666), although this trial was also placed on hold because of possible adverse effects. While findings thus far demonstrate that *ACVR1* over-expression in FOP and control mice is well-tolerated, additional pre-clinical safety studies are required to evaluate whether *ACVR1* over-expression should be considered a possible therapeutic modality. Additionally, of immediate importance will be to establish efficient delivery systems to relevant cell types that minimally activate the host immune system, given the well-known connection between tissue inflammation as both a trigger and manifestation of flare-ups in patients with FOP (*60–62*).

## Materials and Methods

### Generation of *R26^nACVR1^* and *R26^ACVR1^* knock-in mice

*R26^NACVR1^* knockin line was designed to express HA-tagged human ACVR1 (hACVR1-HA) upon Cre-dependent excision of the floxed PGKNeo-stop cassette located between CAG promoter and hACVR1-HA. hACVR1-HA sequence was generated by insertion of a HA linker annealed by the following oligomers, 5’-ACCCATACGACGTACCAGATTACGCTAGTCTCTAGCTCGA-3’ and 5’-GCTAGAGACTAGCGTAATCTGGTACGTCGTATGGGT-3’, between the HincII, which is located at the 3’ end of the ACVR1 coding sequence, and the XhoI sites of pCMV-SPORT6-hACVR1 plasmid (Invitrogen). The R26NALK2 knock-in plasmid, pR26-CNhAHA, was constructed by replacing the nlslacZ-3xSV40pA/FRT/EGFP-bGHpA sequence of pR26-CLNFZG (*24*) with the HA-tagged human ACVR1 followed by a SV40pA. The detail cloning strategy and complete sequence of the plasmids are available on request.

ES cell electroporation and production of chimeric mice were performed by the Center for Mouse Genome Modification (CMGM) at UConn Health. The pR26-CNhAHA plasmid was linearized with Acc65I and electroporated into 129S6/C57BL/6 hybrid ES cells (D1: established by CMGM). Nested PCR was used to screen for homologous recombination on both 5’ and 3’ ends with the following primers; 5’ end: 1^st^ PCR, 5’-TTCCTCTCAATATGCTGCACACAAA-3’ and 5’-GGAACTCCATATATGGGCTATGAACT-3’, 2^nd^ PCR, 5’-AGGCAGGGAAAACGACAAAATCTGG-3’ and 5’-GGAACTCCATATATGGGCTATGAACT-3’; 3’ end: 1^st^ PCR, 5’-TGTGGTTTGTCCAAACTCATCAATGT-3’ and 5’-GATGCCCAATTCCAACTGTGAAGAC-3’, 2^nd^ PCR, 5’-TGTGGTTTGTCCAAACTCATCAATGT-3’ and 5’-TGGTTCACACCACAAATGAACAGTGC-3’. The targeted allele generated 2.4 kb and 5.8 kb diagnostic products with the 5’ and 3’ nested PCR reactions, respectively. Chimeric mice were produced from three targeted ES clones by aggregation with CD1 embryos. Germ line transmission of the targeted allele was assessed by PCR for the floxed PGKNeo-stop cassette/hACVR1 boundary (5’-GATCAGCAGCCTCTGTTCCACA-3’ and 5’-CACGTCTCGGGGATTGAGGC-3’; 653 bp), and the nested PCR for the 5’ and 3’ recombination regions. One *R26^nACVR1^* knockin line was established.

### Mice, crosses, and genotyping

Tie2-Cre transgenic mice were kindly provided by Dr. Tom Sato (UT Southwestern). Pdgfra-Cre transgenic mice (<u>Tg(Pdgfra-cre)1Clc,</u> #013148), *Hprt1^Cre^* (*Hprt^tm1(CAG-cre)Mnn^*, #004302), *Hprt1^iCreER^* (*Hprt^tm345(icre/ERT2)Ems^/Mmjax*, MMRRC stock #037075) and *R26^FLPo^* (*Gt(ROSA)26Sor^tm2(FLP*)Sor^*, #012930) knockin mice were obtained from Jackson Laboratories (USA). *R26^NG^* (*24*), *Alk2^Flox^* (*32, 63*) (referred to here as *Acvr1^flox^*), and *Acvr1^tnR206H^* (*9*) were described previously. *Hprt1^iCre^* was generated by a cross between *Hprt1^iCreER/+^* and *R26^FLPo/+^* mice. *Acvr1^loxP^* was generated by a cross between *Acvr1^flox/+^* and *Hprt1^iCre/+^* mice. *Acvr1^R206H^* and *R26^ACVR1^* were generated by a cross between *Acvr1^tnR206H/+^*;*R26^nACVR1/+^* and *Hprt1^iCre/+^* mice. The *Acvr1^R206H^* allele can be maintained in the presence of *R26^ACVR1^* allele. Transmission of the constitutively recombined *Acvr1^R206H^* allele was verified by the absence of tdTomato red fluorescence and the presence of a 203 bp PCR product using the following primers; 5’-CAACAGGGTTATCTGATGG-3’ and 5’-TCACATGTCCAGAGTTGCT-3’. The following primers were used for genotyping of the other alleles: *Acvr1^tnR206H^* : 5’-GCTAACCATGTTCATGCCTTC-3’ and 5’-AGCGCATGAACTCCTTGATGAC-3’ (144 bp); *Acvr1^flox^*: 5’-TGCTGTCTTTTAACTCCTGGGATC-3’ and 5’-TCTCACCTTCCCTATGACTCTTAG-3’ (0.51 kbp); *Acvr1^loxP^*: 5’-TGCTGTCTTTTAACTCCTGGGATC-3’ and 5’-TCTAAGAGCCATGACAGAGGTTG-3’ (0.6 kbp); *Hprt^iCre^*: 5’-GCTAAAGAGTTGAACGCAAAGGTG-3’ and 5’-GGGCTATGAACTAATGACCCCGTA-3’ (0.6 kbp); Tie2-Cre, Pdgfra-Cre and *Hprt1^Cre^*: 5’-GCGGTCTGGCAGTAAAAACTATC -3’ and 5’-GTTCGAACGCTAGAGCCTGTTT-3’ (174 bp); *R26^ACVR1^*: 5’-GCTAACCATGTTCATGCCTTC-3’ and 5’-CACGTCTCGGGGATTGAGGC-3’ (688 bp) and *R26^NG^*: 5’-GATCAGCAGCCTCTGTTCCACA-3’ and 5’-CGCTGAACTTGTGGCCGTTTAC-3’ (264 bp). Experimental mice were maintained on a mixed background predominantly comprised of FVB, CD1 and C57BL/6 strains.

### Muscle injury by cardiotoxin injection

Muscle injury was performed by injecting 100 µL of 10 µM Naja pallida cardiotoxin (Lotoxan, #L8102) in PBS into the tibialis anterior muscle. HO formation was analyzed at 3 dpi, 8 dpi and 20-24 days post-injury.

### Histology, immunohistochemistry, and µCT imaging

Samples were either directly embedded in Tissue Plus O.C.T. compound (Fisher) immediately after dissection, or fixed in 4% paraformaldehyde/PBS, washed in PBS, and incubated in 30% sucrose in PBS before embedding. For phospho-SMAD1/5/8 immunohistochemical detection, cryosections of paraformaldehyde-fixed samples were subjected to antigen-retrieval as described (*64*). For HRP/DAB-based signal detection, sections were incubated in 0.1% H_2_O_2_/PBS for 30 min. Sections were blocked in 2% BSA/PBS/0.2% Triton X-100 (PBST) supplemented with 50 mM glycine with primary antibodies in PBST supplemented with 10 mM glycine. A mixture of rabbit anti-pSmad1/5/9 (D5B10) and anti-pSMAD1/5 (41D10) monoclonal antibodies (Cell Signaling, #13820 and #9516, respectively), diluted 1:100 each, was used for pSMAD1/5/8 detection. Rabbit anti-SOX9 antibody (Millipore, #AB5535) and rabbit anti-HA antibody (Zymed, #71-5500) were used at 1:1000 and 1:50 dilutions, respectively. Biotin-XX-goat anti-rabbit IgG (Thermo Fisher, #B2770) was used as a secondary antibody at 1:2000 dilution in PBS/5% goat serum/0.2% Triton X-100. Alexa Fluor 647-streptavidin (Thermo Fisher, #S21374) was used for fluorescent detection of the target proteins. Vectorstain Elite ABC-HRP kit (Vector Lab, #PK-6100) and Metal Enhanced DAB Substrate kit (Thermo Fisher, #34065) were used for chromogenic detection of the target proteins.

Alcian blue/Alizarin red whole mount skeletal staining (*65*) and µCT imaging (*9*) were performed as described (*9*).

### Cell isolation and culturing for *Acvr1* mRNA and protein quantification

Total hindlimb muscle tissue was dissected from adult mice, physically dissociated into small pieces, and incubated in 0.2% collagenase type 1/DMEM/1X Penicillin-streptomycin (Pen-Strep, Caisson, #PSL01) for 90 min at 37°C with gentle shaking. Digestion was terminated by addition of FBS to a final concentration of 10%. After filtering through a polyester fiber filter, the lysate was centrifuged at 800 x g for 2 min. The cells were subjected to a red-blood cell lysis, filtered through a 100 µm cell strainer (Falcon, #352360) and centrifuged at 300 x g for 5 min. After counting with a hemocytometer, cells were centrifuged at 800 x g for 2 min and resuspended in DMEM/10% FBS/1X Pen-Strep.

2.5 X 10^6^ cells were plated in a 6 cm tissue culture dish and cultured for 90 min in DMEM/10% FBS/1X Pen-Strep. After removing non-adherent cells by flushing with PBS, adherent cells were further cultured overnight in DMEM/10% DMEM/1X Pen-Strep. After rinsing with PBS, cells were processed for *Acvr1* mRNA or protein quantification.

### *Acvr1* mRNA quantification

RNA from the adherent cells was extracted using QIAGEN RNeasy Micro kit according to the manufacturer’s manual. RNA from 14.5 dpc whole fetuses was prepared using QIAGEN RNeasy Midi kit. Complementary DNA (cDNA) was synthesized using BioRad iScript Advanced cDNA Synthesis Kit (#1725038) according to the manufacturer’s instructions. Absolute quantification of target transcripts was determined by BioRad QX200 Droplet Digital PCR system using QX200 ddPCR EvaGreen Supermix (#1864033) according to the manufacturer’s instructions. The following PCR primer sets were used for each gene transcripts; *Acvr1* wild type: 5’-CAGCACTCTAGCGGAACTAC-3’ and 5’-CTCCAACAGGGTTATCTGGC-3’ (106-bp), *Acvr1^R206H^*: 5’-CAGCACTCTAGCGGAACTAC-3’ and 5’-CAACAGGGTTATCTGATGG-3’ (103-bp), *β Actin*: 5’-TAGGCACCAGGGTGTGATGG-3’ and 5’-GTACATGGCTGGGGTGTTGAA-3’ (286-bp), *Pcna*: 5’-TGGAGAGCTTGGCAATGGGAA-3’ and 5’-GCAAACGTTAGGTGAACAGGCT-3’ (108-bp), *R26^ACVR1^*: 5’-AATCCATCCGCAAGACTCAC-3’ and 5’-CTCGAGCTAGAGACTAGCGT-3’ (129-bp). After droplet generation with the QX200 Droplet Generator (BioRad), thermal cycling was performed with a C1000 Thermal Cycler (BioRad) as follows: 95 °C for 5 min, 40 cycles of 95 °C for 30 sec and 60 °C for 60 sec, followed by one each of 4°C for 5 min and 98 °C for 10 min, all at a ramp rate of 2°C per second. The droplets were subsequently read by a QX200 Droplet Reader and the data were analyzed with QuantaSoft v.1.7.4 software (Bio-Rad).

### Western blotting for detection of ACVR1

Adherent cells were collected in 100 µL 2X RIPA buffer supplemented with 2 mM PMSF. Total volume of cell lysate was adjusted to 200 µL with PBS. 14.5 dpc fetus was homogenized in 4 ml of 1X RIPA buffer supplemented with 1 mM PMSF, rocked for 2 hours at 4°C, centrifuged at 10,000 rpm for 20 min at 4°C, and the supernatant collected. After protein quantification, an equal volume of 2X Laemmli SDS-sample buffer was added to the protein lysate. A BioRad Mini Trans-Blot system was used to perform SDS-PAGE and protein transfer onto PVDF membrane according to the manufacturer’s instructions. Immunoblotting was performed using SuperSignal West Pico PLUS (Thermo Fisher, #34577) according to the manufacturer’s instructions. For sequential immunodetection of multiple epitopes, stripping was performed by incubating used membrane in 62.5 mM Tris-HCl pH 6.8/ 1% SDS/ 100 mM β-mercaptoethanol for 40 min at 50°C followed by rinsing in water. The following antibodies and dilutions were used: rabbit monoclonal anti-ACVR1 antibody (Abcam, #155981) diluted 1:1000, rabbit anti-PCNA antibody (Santa Cruz, #FL-261) diluted 1:1000, mouse monoclonal anti-HA antibody (2-2.2.14, Thermo Fisher, #26183) diluted 1:10^4^, rabbit anti-mouse IgG (Alexa Fluor 647-conjugated, Thermo Fisher, #A21239) diluted 1:10^4^ and HRP-conjugated goat anti-rabbit IgG (Cell Signaling, #7074) diluted 1:10^4^. Chemiluminescence was detected using FluorChem HD2 (ProteinSimple) or IVIS SpectrumCT (Perkin Elmer) instruments.

### Cell isolation, culturing, and Western blotting for pSMAD1/5/8

Total hindlimb skeletal muscle was harvested from mice of the indicated genotypes, minced, enzymatically digested, and filtered as previously described (*9*). The resulting crude mixture of cells from each mouse was seeded into a 100 mm cell culture dish (NEST, #704004) and grown to 60-80% confluence, over 3-4 days, in growth media containing 20% fetal bovine serum (FBS) (R&D Systems, #S11150) and 1% Pen/Strep (Sigma, P4333) in Dulbecco’s Modified Eagle Medium (DMEM) (Life Technologies, #11995-073). Crude plastic-adherent cells were then dissociated from the culture dishes using Accumax (Innovative Cell Technologies, AM105), blocked with 10% FBS in PBS, and stained for 30 minutes on ice with the following fluorescently-conjugated antibodies at indicated dilutions: anti-PDGFRα-APC (1:100; eBioscience, #17-1401-81), anti-SCA-1-v450 (1:400; BD Bioscience#560653), anti-CD31-bv711 (1:500; BD Biosciences, #740690), and anti-CD45-bv711 (1:500; BD Biosciences, #563709). 7-AAD was also added to cell suspensions at 0.5µg/mL 10 minutes prior to identify and exclude dead/dying cells. Using a FACSAria II flow cytometer (BD Biosciences), control FAPs, with and without the *R26^ACVR1^* allele, were isolated as live single cells expressing the cell surface profile, CD31-CD45-SCA-1+PDGFRα+. R206H-FAPs, with and without the *R26^ACVR1^* allele, were identified with the additional selection criteria tdTomato-GFP+, which selects for only *Acvr1^tnR206H^*-recombined FAPs.

FACS-isolated FAPs were seeded in wells of 6-well plates at 2,000 cells/cm^2^ in growth media. After reaching near-confluence, FAPs were serum starved in DMEM for 2 hours before a 1-hour incubation with either activin A or BMP6 at the indicated concentrations. FAPs were washed with cold PBS, scraped off the dish in cold RIPA lysis buffer, and vortexed to complete lysis. Total protein concentration was measured using the DC protein assay (Bio-Rad), and equal amounts of total protein were loaded per lane of the same gel for SDS-PAGE. After the transfer step, PVDF membranes were blocked in 5% non-fat milk in PBS with Tween-20, and incubated with rabbit anti-pSMAD1/5/8 (1:1,000; CST, #13820) and mouse anti-β-actin (1:2,500; CST #3700) overnight at 4°C. Goat anti-rabbit IgG HRP-linked antibody (CST, #7074) was used either as a secondary antibody to detect pSMAD1/5/8, or as a tertiary antibody after incubation with rabbit anti-mouse IgG Alexa Fluor 647 to detect β-actin. Membranes were incubated in SuperSignal™ West Pico PLUS Chemiluminescent Substrate (Thermo Fisher Scientific) and imaged using the IVIS SpectrumCT (Perkin-Elmer). pSMAD1/5/8 and β-actin bands were imaged separately, as separate secondary antibody incubations of the same membrane were required. Blots were quantified using ImageJ, wherein the integrated density values (arbitrary units) of pSMAD1/5/8 bands were divided by those of β-actin bands in the same sample to generate normalized integrated density values. The normalized integrated density values from individual, representative blots are displayed as bar graphs, which were generated in GraphPad Prism v7.05.

## List of Supplementary Materials

**Figure S1.** *Acvr1^tnR206H^* and *R26^nACVR1^* conditional alleles.

**Figure S2.** Limb skeletal defects in *Acvr1^R206H/+^* mutant mice.

**Figure S3.** Effects of *Acvr1^R206H^* and *R26^ACVR1^* alleles on rib development.

**Figure S4.** Spontaneous skeletal expansions in the legs of some aged mice that carried the *R26^ACVR1^* allele.

**Figure S5.** Inhibition of exacerbated HO by *ACVR1* over-expression.

**Figure S6.** Injury-induced hyper-cellularity of intermuscular fascia.

**Figure S7.** No discernible effect of *ACVR1* over-expression on pSMAD1/5/8 or SOX9 levels in the TA muscle 3 days post-injury.

## Acknowledgments

We thank Dr. Vesa Kaartinen for *Acvr1* knockout mice, Acceleron Pharma for the anti-activin A antibody, Dr. Wu He, Director of the Flow Cytometry Facility, for technical assistance with FACS, and members of the Goldhamer Lab for helpful discussions throughout the course of this work. This work was funded by NIH grants R01AR072052 and R21AR074584 to D.J.G. The authors declare no competing interests.

## Author contributions

D.J.G., M.Y., and S.J.S. designed the study. M.Y., S.J.S., and S.Y. performed the experiments. D.J.G., M.Y., and S.J.S. analyzed and interpreted the data. D.J.G. wrote the manuscript with essential input from M.Y. and S.J.S.

## Notes

### Competing Interest Statement

The authors have declared no competing interest.

