## Supplementary Figures 1-7 for "Over-expression of wild-type *ACVR1* in fibrodysplasia ossificans progressiva mice rescues perinatal lethality and inhibits heterotopic ossification"

### Supplementary Materials

Supplementary Figures 1-7.

**Supplementary Figure 1**

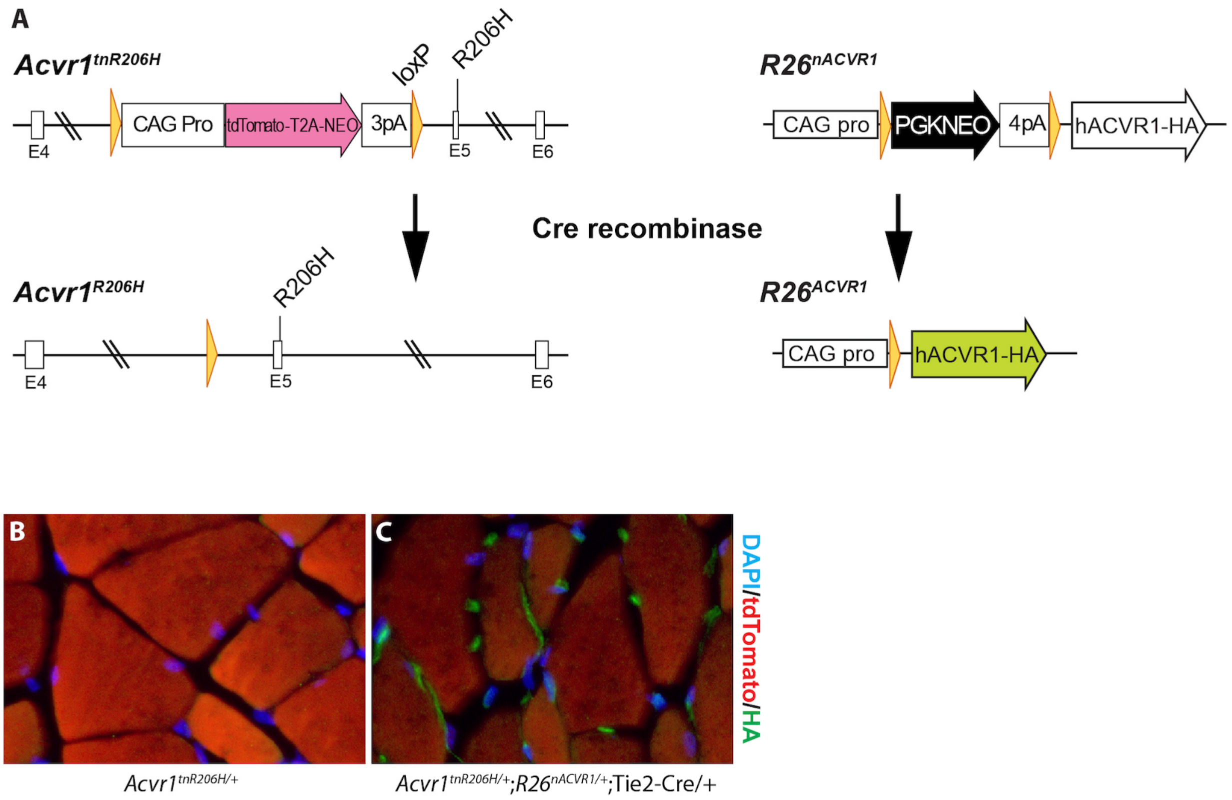


**Figure S1**. *Acvr1*$^{tnR206H}$ and *R26*$^{nACVR1}$ conditional alleles. **(A)** Schematic representation of Cre-dependent recombination of *Acvr1*$^{tnR206H}$ and *R26*$^{nACVR1}$ alleles. The *Acvr1*$^{tnR206H}$ allele is described in ref. *(9)*. **(B, C)** Detection of HA-tagged human ACVR1 protein by HA-immunofluorescence. **(B)** Fluorescent image of a cross-section of the TA muscle of an *Acvr1*$^{tnR206H/+}$ control mouse. The unrecombined *Acvr1*$^{tnR206H}$ allele expresses tdTomato and labels the myofibers. **(C)** Muscle section from an *Acvr1*$^{tnR206H/+}$;*R26*$^{nACVR1/+}$;Tie2-Cre mouse. Vascular elements and Tie2+ mesenchymal progenitors express ACVR1-HA (green).

**Supplementary Figure 2**

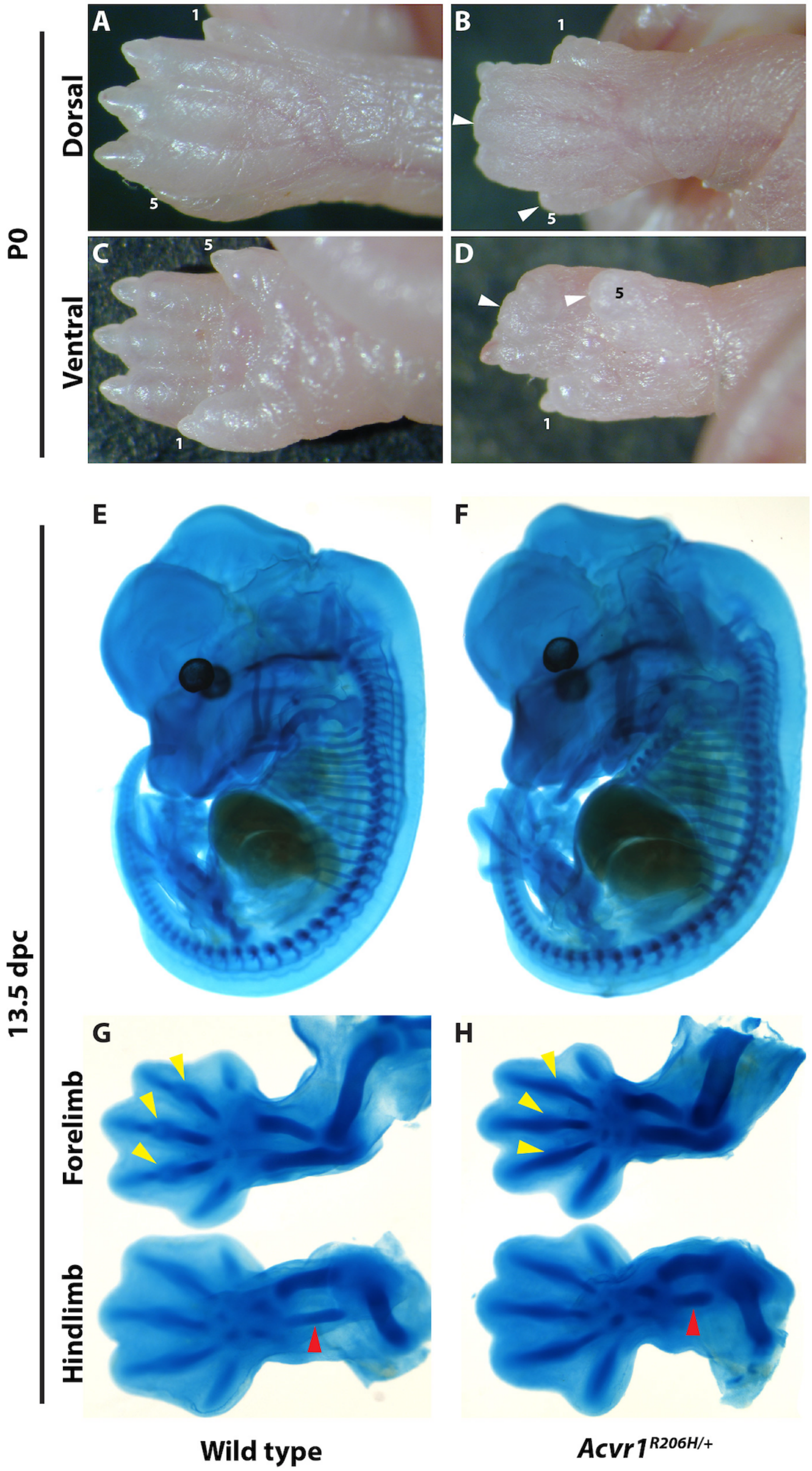

**Figure S2**. Limb skeletal defects in *Acvr1*$^{R206H/+}$ mutant mice. **(A-D)** Defects of the hindlimb autopod in P0 mice. The first and fifth toes are labeled 1 and 5, respectively. In *Acvr1*$^{R206H/+}$ neonates, separation of the toes is incomplete, and some toes are distally truncated (white arrowheads). **(E-H)** Abnormal limb skeletal patterning in *Acvr1*$^{R206H/+}$ embryos. Alcian blue staining of wild-type (E, G) and *Acvr1*$^{R206H/+}$ (F, H) whole mount embryos and limbs at 13.5 dpc. Early manifestations of skeletal defects in *Acvr1*$^{R206H/+}$ animals include an apparent defect in formation of the metacarpophalangeal joints in the forelimb (yellow arrowheads), and shortening of the fibula (red arrowheads).

**Supplementary Figure 3**

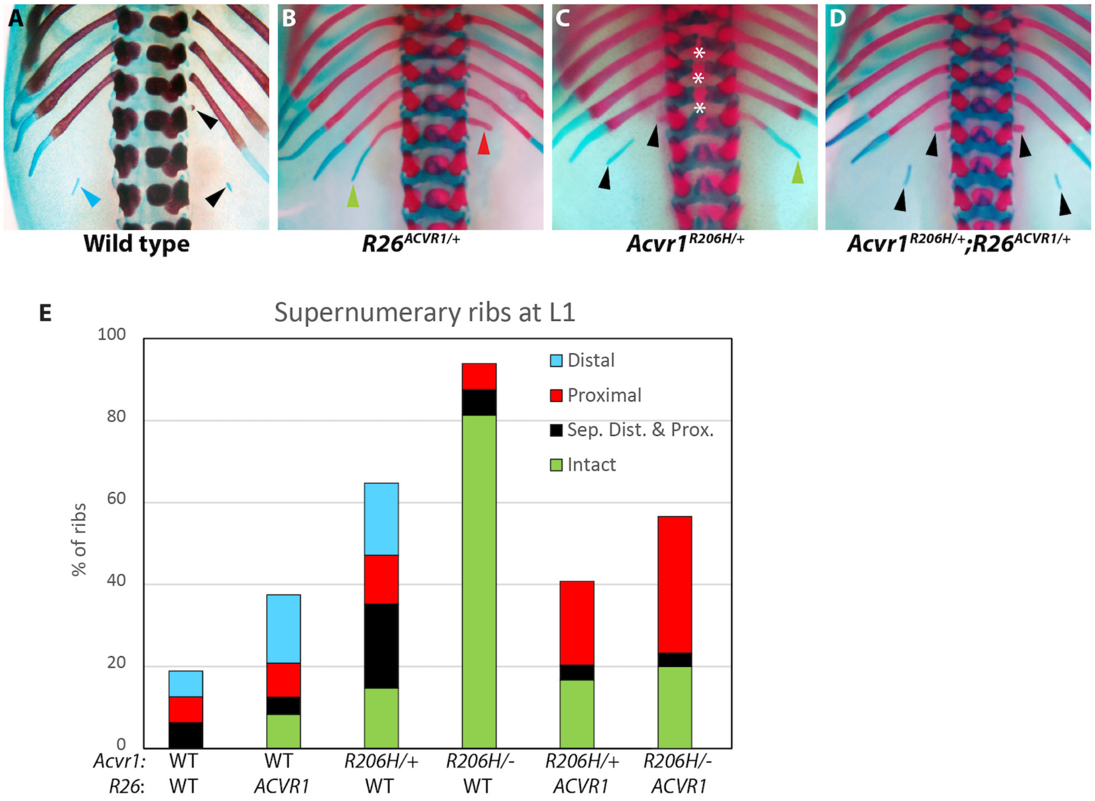


**Figure S3**. Effects of *Acvr1*$^{R206H}$ and *R26*$^{ACVR1}$ alleles on rib development. **(A-D)** Representative images of P0 neonates with supernumerary ribs at the 1st lumbar (L1) level. **(A)** wild-type (WT) neonate with a floating distal cartilage (blue arrowheads) on the left side and a separated proximal protrusion and floating cartilage (black arrowheads) on the right side. **(B)** *R26*$^{ACVR1/+}$ mouse with an intact L1 rib (left, green arrowhead) and a proximal bony protrusion (right, red arrowhead). **(C)** *Acvr1*$^{R206H/+}$ mouse with a proximal protrusion and a floating cartilage (left, black arrowheads) and an intact extra rib (right, green arrowhead). Area of mild spina bifida at the lower-thoracic/upper-lumbar level is denoted with asterisks. **(D)** *Acvr1*$^{R206H/+}$;*R26*$^{ACVR1/+}$ mouse with a pair of separated proximal and distal rib compartments on both left and right L1 sides. **(E)** Quantification of the occurrence of L1 supernumerary ribs with different developmental structures. Left- and right-side structures were counted separately.

**Supplementary Figure 4**

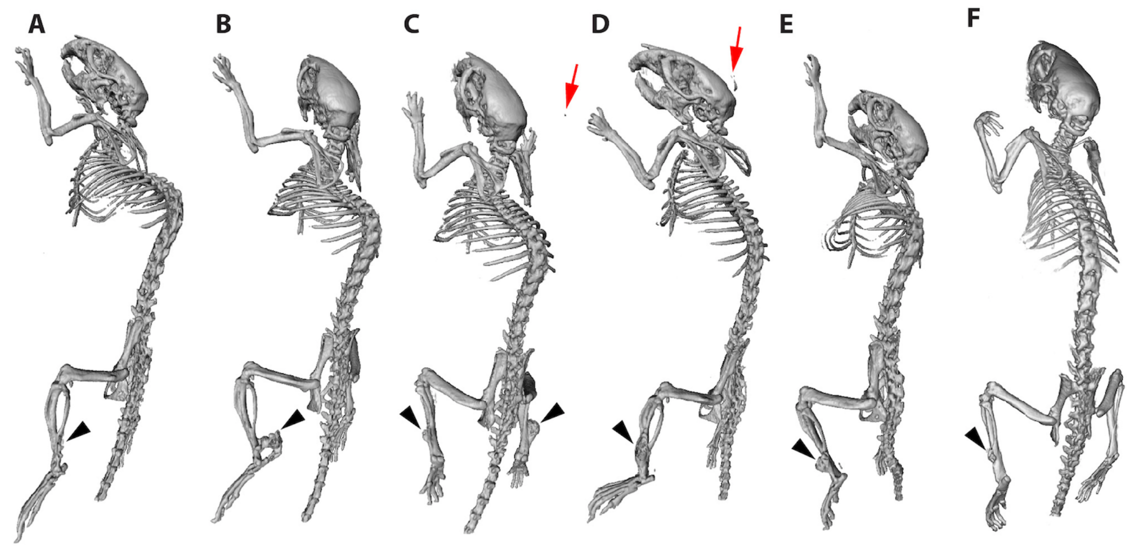


| *Acvr1* | *R206H/+* | *R206H/+* | *R206H/+* | *R206H/+* | *R206H/+* | *tnR206H/+* |
|---|---|---|---|---|---|---|
| *R26* | *ACVR1/+* | *ACVR1/+* | *ACVR1/+* | *ACVR1/+* | *ACVR1/+* | *ACVR1/+* |
| Cre type | None | None | None | *Hprt1*$^{iCre}$ | None | Pdgfra-Cre |
| Age | 15.8 MO | 13.1 MO | 15.7 MO | 15.3 MO | 13.8 MO | 10.3 MO |

**Figure S4**. Spontaneous skeletal expansions in the legs of some aged FOP mice that carried the *R26*$^{ACVR1}$ allele. The table shows the genotype and age (months) of each mouse. **(A - E)** µCT images of all five *Acvr1*$^{R206H/+}$;*R26*$^{ACVR1/+}$ mice (n=17) that developed abnormal skeletal expansions, which presented at the distal tibia (black arrowheads). All affected mice were females. In 6 of 17 mice, HO developed in the ear pierced with a metal tag (red arrows in C and D); this HO depended on *Acvr1*$^{R206H}$ but was not specific to aged mice. **(F)** *Acvr1*$^{tnR206H/+}$;*R26*$^{ACVR1/+}$;Pdgfra-Cre mouse with an aberrant skeletal growth on the distal tibia (black arrowhead).

**Supplementary Figure 5**

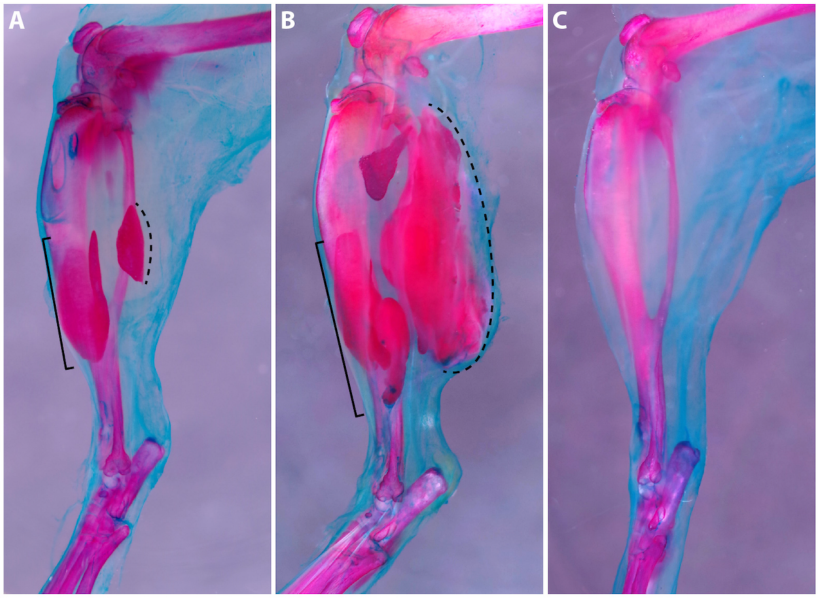


| *Acvr1* | *tnR206H/+* | *tnR206H/flox* | *tnR206H/flox* |
|---|---|---|---|
| *R26* | +/+ | +/+ | *nACVR1/+* |
| Cre type | Tie2-Cre/+ | Tie2-Cre/+ | Tie2-Cre/+ |

**Figure S5**. Inhibition of exacerbated HO by *ACVR1* over-expression. The table shows the genotype of each mouse. **(A)** Representative injury-induced HO response of adult *Acvr1*$^{tnR206H/+}$;Tie2-Cre mouse. The limb was ABAR-stained 21-day after cardiotoxin injection of the TA muscle. **(B)** Removal of the wild-type *Acvr1* allele exacerbated HO. Hatched lines and brackets in (A) and (B) indicate heterotopic bone in the soft tissue and aberrant skeletal expansion on the surface of the tibia, respectively. **(C)** Over-expression of *ACVR1* completely prevented HO in *Acvr1*$^{tnR206H/flox}$;*R26*$^{ACVR1/+}$;Tie2-Cre mice.

**Supplementary Figure 6**

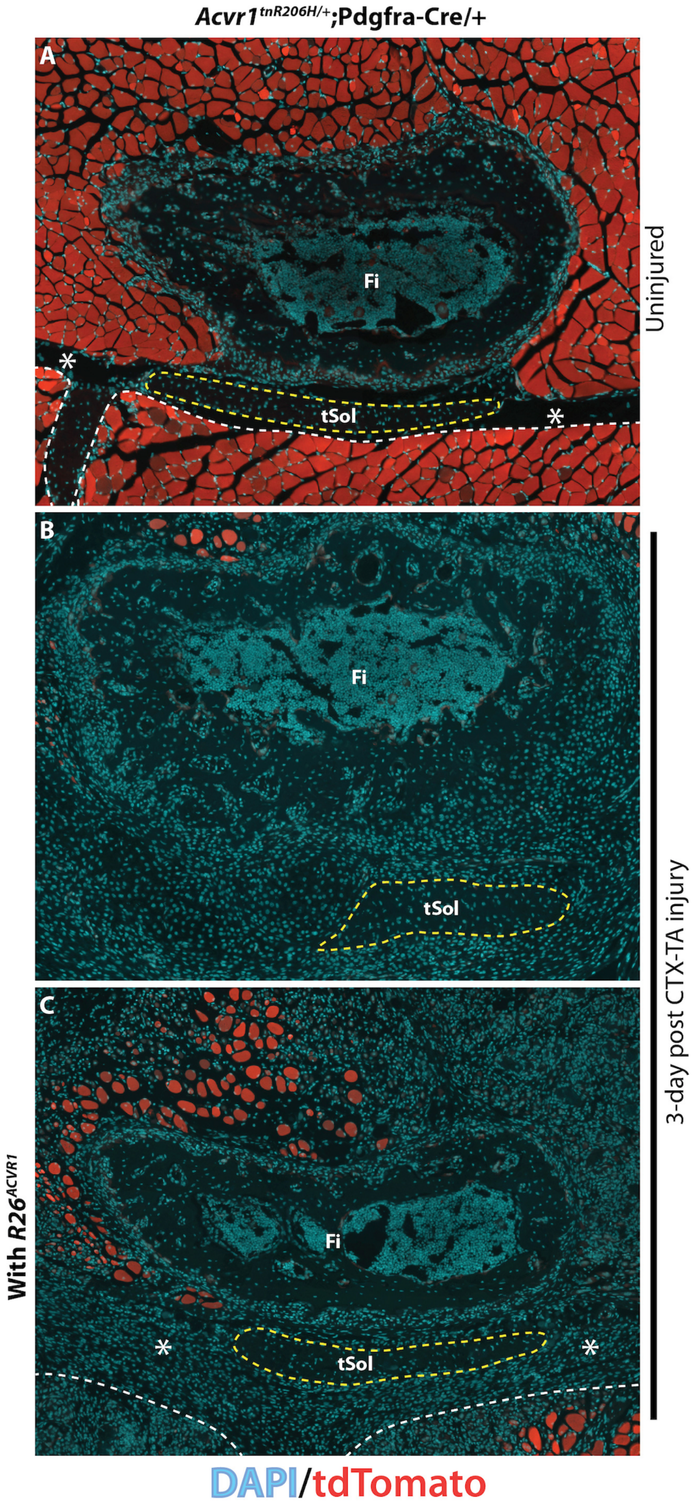

**Figure S6**. Injury-induced hyper-cellularity of intermuscular fascia. Tissue was collected 3 days after cardiotoxin-induced injury of the tibialis anterior (TA) muscle. **(A)** Transverse section through the lower limb showing the typical appearance of the transverse intermuscular septum (asterisks). Note the low density of mononuclear cells in the septum. **(B)** In FOP mice, the transverse septum has lost its well-defined structure and become dramatically hypercellularized by 3 days post-injury. **(C)** The response to muscle injury of FOP mice carrying the $R26^{ACVR1}$ allele is comparable to that of control mice. Asterisks denote the widened septum. White dashed lines represent the boundary between the septum and adjacent skeletal muscle. Muscle fibers express tdTomato from the unrecombined $Acvr1^{tnR206H}$ allele (red). Red fluorescence of unrecombined mononuclear cells is not evident, particularly at this magnification, because the exposure time was set to optimally capture the intense fluorescence of the muscle fibers. Fi, fibula; tSol, soleus tendon.

**Supplementary Figure 7**

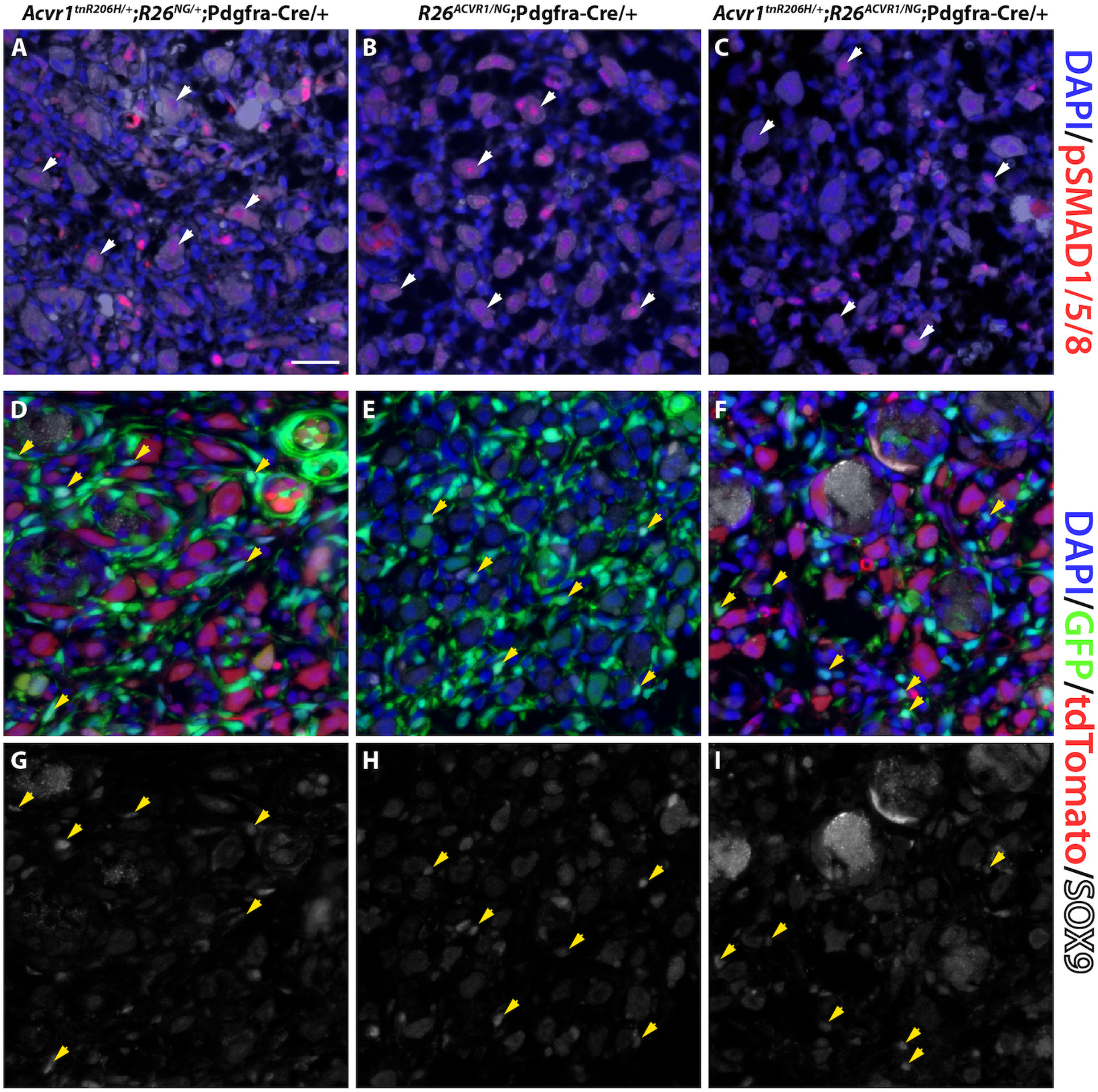

**Figure S7**. No discernible effect of *ACVR1* over-expression on pSMAD1/5/8 or SOX9 levels in the TA muscle 3 days post-injury. **(A-C)** The nuclei of some nascent muscle fibers are weakly positive for pSMAD1/5/8 irrespective of genotype. Autofluorescence through the green channel was colorized white to show the tissue structure. Note that GFP fluorescence from the recombined $R26^{NG}$ reporter was destroyed by the antigen retrieval method used to detect pSMAD1/5/8. White arrowheads indicate representative regenerating myofibers with pSMAD1/5/8-positive nuclei. **(D-F)** Simultaneous detection of GFP from the recombined $R26^{NG}$ reporter, tdTomato from the unrecombined $Acvr1^{tnR206H}$ allele, and DAPI. Nascent muscle fibers are the source of most of the tdTomato signal in (D) and (F). **(G-I)** Same sections as in (D-F) showing staining for SOX9. The yellow arrowheads in (D-I) indicate representative GFP+ recombined cells with weak SOX9 staining.
